# A molecular clock controls periodically driven cell migration in confined spaces

**DOI:** 10.1101/2020.12.29.424673

**Authors:** Sung Hoon Lee, Archer Hamidzadeh, M. Sulaiman Yousafzai, Visar Ajeti, Hao Chang, Michael Murrell, Andre Levchenko

**Affiliations:** Yale Systems Biology Institute, Yale University, West Haven, CT 06516, USA; Department of Biomedical Engineering, Yale University, New Haven, CT 06520, USA; Department of Physics, Yale University, New Haven, CT 06520, USA; Department of Biomedical Engineering, University of Connecticut Health Center, Farmington, CT 06032, USA

## Abstract

Navigation through dense, physically confining extracellular matrix is common in invasive cell spread and tissue re-organization, but is still poorly understood. Here, we show that this migration is mediated by cyclic changes in the activity of a small GTP-ase RhoA, dependent on the oscillatory changes in the activity and abundance of the RhoA Guanine Exchange Factor, GEF-H1, triggered by a persistent increase in the intracellular Ca^2+^ levels. We show that the molecular clock driving these cyclic changes is mediated by two coupled negative feedback loops, dependent on the microtubule dynamics, with the frequency that can be experimentally modulated based on a predictive mathematical model. We further demonstrate that an increasing frequency of the clock translates into a faster cell migration within physically confining spaces. This work lays the foundation for a better understanding of the molecular mechanisms dynamically driving cell migration in complex environments.

## Introduction

Cell migration is a highly regulated process relying on complex reorganization of intracellular cytoskeleton, controlled by a large variety of extracellular cues and intracellular regulatory networks. Migrating cells can employ diverse modes of locomotion in different micro-environments (Friedl and Wolf, 2010). In particular, during the so-called 3D migration in dense extracellular matrices (ECM), multiple cell types can switch to amoeboid movement, characterized by a weak adhesion to ECM and a Myosin II-dependent motility (Liu et al., 2015). This mode of migration is frequently contrasted with 1D migration and more commonly analyzed 2D migration, which can also occur *in vivo.* 1D migration can be triggered and sustained in sparser ECM, with cells moving along a single ECM fiber or multiple aligned fibers (Doyle et al., 2009; Kim et al., 2009). 2D migration may occur along very dense matrix structures, such as basement membranes, and very commonly in lab experiments, including cell migration in most culture conditions (Abrams et al., 2002; Doyle et al., 2013; Kim et al., 2012). The 3D migration mode is distinct from these two and is particularly important as it can mediate the key processes accompanying a highly invasive spread of cells into dense surrounding ECM, occurring in such key physiological processes as the extension of a growing neovascular sprouts led by a migrating Tip cell or invasion of trophoblasts during formation of placenta (Geudens and Gerhardt, 2011; Knofler and Pollheimer, 2013). It also can underlie dissemination of multiple types of cancer cells (Friedl and Alexander, 2011). Unfortunately, compared to 2D migration, regulation of 3D migration in highly confined physical spaces is still poorly understood, prompting the need for new tools and methods for its analysis.

Cell migration relies on a careful orchestration of complex molecular processes in space and time, particularly since they may lead to ostensibly contradictory outcomes. For instance, actin polymerization and ensuing formation and expansion of cellular projections, such as pseudopods and filopodia, needs to be carefully balanced with the contraction of cortical cytoskeleton and propulsive force generation (Ridley et al., 2003). In space, the coordination of cytoskeletal rearrangements is achieved through a complex signaling crosstalk mediated by small GTPases of the Rho family, mediating spatial polarization of actin polymerization and contraction (Chauhan et al., 2011; Lawson and Ridley, 2018; Park et al., 2017). However, it is less clear if and how these processes may be coordinated in time. Indeed, it is frequently thought that cell migration is cyclic in nature, with the extension of the cell front and contraction of its rear alternating in time during the locomotion process (Devreotes and Horwitz, 2015; Petrie et al., 2009). However, it is not clear how this oscillatory migratory dynamics is driven and how the molecular clock activities might operate in diverse migratory modes.

Cell migration depends on dynamic reorganization of the cytoskeleton. Whereas the role of actin dynamics in controlling cell motility is frequently at the focus on analysis, the role of microtubule cytoskeleton is relatively poorly characterized. Historically, microtubules have been treated as accessory structures enabling e.g., the transport of key cargo needed for maintenance of cell polarity and migration (Gundersen and Bulinski, 1988; Hirokawa et al., 1998; Siegrist and Doe, 2007). However, more recently, the integrity and stability of microtubule cytoskeleton have been seen as more immediate regulators of dynamic changes in cell shape and motility (Belotti et al., 1996; Bijman et al., 2006; Dogterom and Koenderink, 2019; Yang et al., 2010). Pharmacological perturbations of microtubules, motivated in part by the current chemotherapeutic modalities, revealed that microtubule stabilization can inhibit cell migration (Prahl et al., 2018). Microtubule destabilization can promote complex dynamic changes in cell shape, including oscillatory changes of cell polarity (Park et al., 2017). Furthermore, these perturbations can have a direct effect on actin cytoskeleton and cell adhesion (Tabdanov et al., 2018). In the context of 3D cell migration, dynamic changes in microtubule cytoskeleton in the rear of migrating cells (the uropod), which is also the site of actin-myosin interactions leading to generation of propulsive forces, are essential for effective cell motility (Bouchet and Akhmanova, 2017; Hind et al., 2016). These findings raise the question of how the dynamic changes of microtubule and actin cytoskeleton are coordinated during cell locomotion, particularly in 3D.

Although, as noted above, many facets of 3D cell migration in confined spaces remain poorly understood, recent research began to reveal important correlates of this process. In particular, it has been observed that mechanical confinement can trigger activation of mechano-sensitive Ca^2+^ channels, including Piezo1/2, resulting in a persistent increase in intracellular Ca^2+^ concentration (Hung et al., 2016; Miyamoto et al., 2014). Ca^2+^ is a pleiotropic second messenger that can also be controlled by other environmental and biochemical stimuli. Thus, unraveling of its putative role in controlling cell migration can help understand how diverse stimuli may impact the motility apparatus. Recently, the effect of intracellular Ca^2+^ on Myosin Light Chain Kinase was implicated as a key determinant of endothelial cell migration *in vitro* and *in vivo* (Noren et al., 2016; Yokota et al., 2015) and the effect on the small GTP-ase RhoA was connected to breast cancer cell migration in confined spaces (Pardo-Pastor et al., 2018). Furthermore, Piezo2 mediated Ca^2+^ signaling was connected to Focal Adhesion Kinase signaling in glioblastoma invasion (Chen et al., 2018). However, given its pleiotropic nature, many other Ca^2+^ effectors may play critical roles in controlling cell migration and coordinating the underlying dynamical molecular processes.

In this study, we explored the complex interplay between Ca^2+^ signaling, microtubule cytoskeleton organization, the activity of small GTPases and the resulting cell migration. We find that this interplay is tightly coordinated in time through two interlinked negative feedback loops regulating the activity and abundance of a GEF protein controlling the activity of RhoA, GEF-H1. This feedback regulation leads to an oscillatory activity of RhoA occurring in synchrony with the microtubule polymerization-depolymerization cycles, generating a molecular clock controlling the speed of cell locomotion in confining environments. The clock can be triggered and sustained by an increase in intracellular Ca^2+^ concentration in response to either spatial confinement or pharmacological perturbations. This analysis reveals a complex and tightly dynamically controlled timer of cell locomotion essential for effective cell locomotion that can operate in diverse cell types and invasive cell processes.

## Results

### Cell migration in physically confined spaces is driven by a molecular clock controlling Ca^2+^ stimulated oscillatory RhoA activity

To explore the relationship between Ca^2+^ regulation and cell motility in mechanically confining 3D microenvironments, we used fluorescent biosensors of Ca^2+^ signaling and of the activity of a small GTPase, RhoA, considered to be a key regulator of this type of cell migration (Hung et al., 2013; Liu et al., 2015). In these experiments, we studied single and collective cell migration in narrow (≤ 6 μm) and shallow (≤ 10 μm) micro-fabricated channels (**Supplementary Figure 1A**) as well as invasive cell migration in 3D collagen matrices (**Figure S1 in Methods S1**). As expected, we found that Ca^2+^ becomes persistently elevated under all these confining conditions within single and collectively migrating cells of the mouse pancreatic endothelial cell line, MS1, expressing a genetically encoded Ca^2+^ biosensor GCaMP5 (**Figure 1A and Figure S3 in Methods S1**) and cells of a human melanoma cell line A375, loaded with the Fluo4 Ca^2+^- sensitive dye (**Supplementary Figure 1B**). Using a live cell RhoA activity reporter (the RhoA2G) in MS1 and A375 cells, we found that RhoA activity oscillated in both cell types (**Figure 1B,C, Supplementary Figure 1C and Methods S2**). Furthermore, cell migration was periodically driven, with peaks of the migratory speed coinciding in A375 cells with the peaks of RhoA activity oscillations (**Figure 1C**). Importantly, in spite of a large variability of the oscillation frequency across individual cells, the frequency of oscillations of RhoA and cell migration were highly correlated (**Figure 1D**). Furthermore, the frequency of the RhoA oscillations and the average RhoA activity during single and collective migration of MS1 cells in confining channels and single cell migration of A375 cells were strongly downregulated in the presence of BAPTA (1,2-bis(o-aminophenoxy)ethane-N,N,N’,N’-tetra-acetic acid) (**Figures 1E,F**), a Ca^2+^-specific chelator, revealing that Ca^2+^ is a major regulator of the oscillatory RhoA dynamics. These results suggested that in cells migrating under spatial confinement, both RhoA dynamics and migration speed can display correlated oscillatory dynamics that may strongly depend on the mechanically induced increase in the intracellular Ca^2+^ concentration.

**Figure 1.**
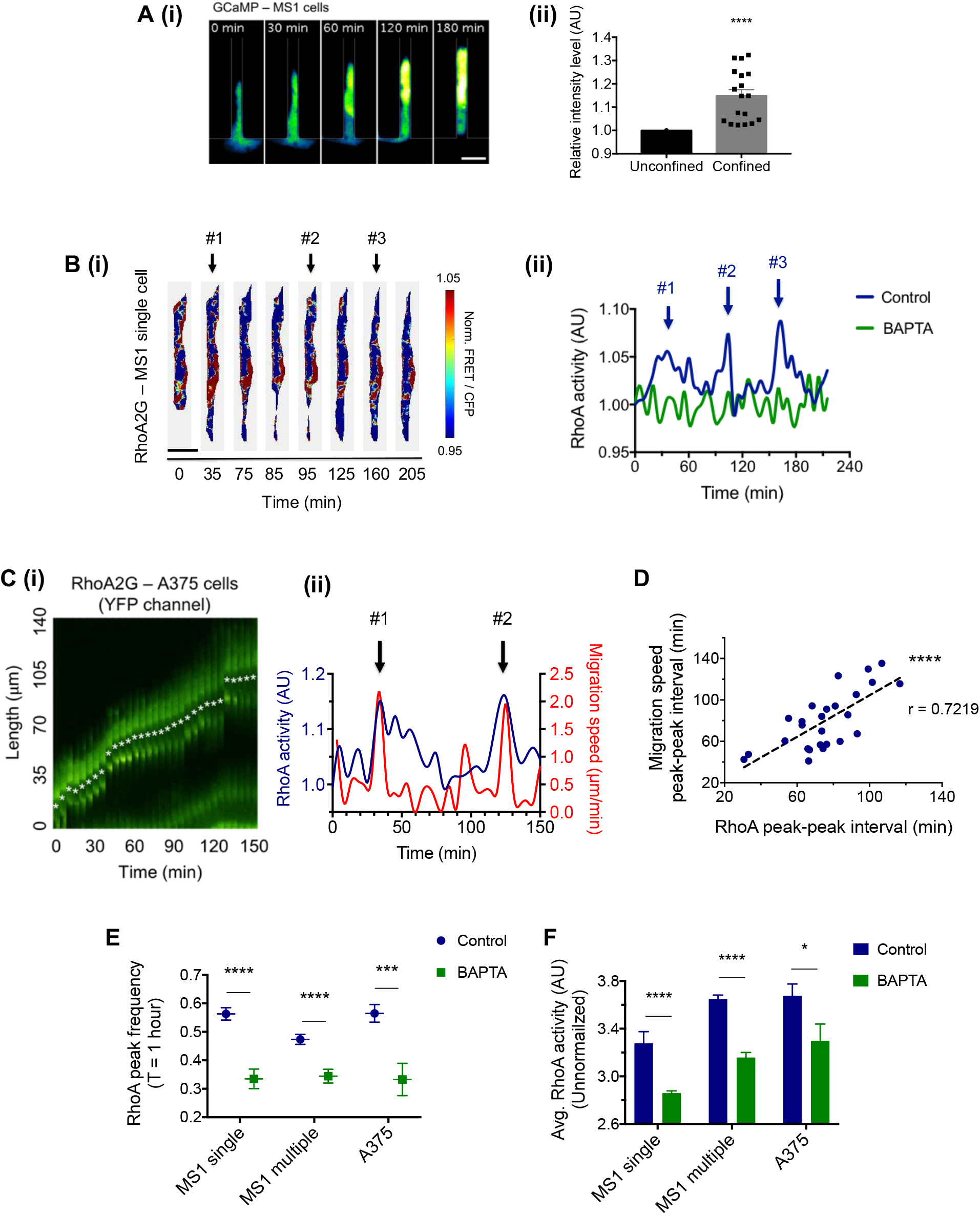
An increase of intracellular Ca^2+^ induced during migration under physical confinement leads to oscillatory RhoA dynamics and cell migration. **(A)** (i) Time-lapse images of GCaMP – MS1 cells migrating in a spatially confining channel (version2). Scale bar: 20 μm. (ii) Comparison of the fluorescence intensity of cells before (unconfined) and after (confined) entering the channel. Data were analyzed by unpaired two-tailed t test with error bar representing s.e.m. (n = 17 cells, **** p < 0.0001) **(B)** (i) Representative time-lapse images of RhoA activity of a single MS1 cell under confinements. Pseudo-color images show the FRET ratio (FRET/CFP) using the RhoA2G biosensor. Scale bar: 20 μm. (ii) RhoA activity of a single MS1 cell migrating in the channel with or without BAPTA (1 μM). Here and elsewhere, the peaks of activity were detected by an algorithm described in the Methods and Supplementary Methods File 2. **(C)** (i) Time-lapse kymograph tracking a A375 cell with the RhoA2G biosensor migrating in the channel (version1). (ii) RhoA activity of the cell shown in (i) superimposed on migration speed plotted over time. The numbers indicate peaks that were detected as described above. **(D)** Scatter plot of RhoA activity peak-peak interval and the corresponding migration speed peak-peak interval of A375 cells. (n=26) **(E)** Comparison of RhoA peak frequency measured over an hour with or without BAPTA (1 μM) for single, multiple MS1 cells and single A375 cells under physical confinements (in channels: version1 for A375 cells, version2 for MS1 cells). Data were analyzed by two-way ANOVA. (n = 60 cells (control, MS1 single), n = 38 cells (BAPTA, MS1 single), n = 70 groups of cells (control, MS1 multiple) and n = 58 groups of cells (BAPTA, MS1 multiple), n = 18 cells (control, A375), n = 11 cells (BAPTA, A375) *** P = 0.0009, **** P < 0.0001) **(F)** Comparison of averaged RhoA activity for single, multiple MS1 cells and A375 cells under confinements. Data were analyzed by Two-way ANOVA. (n = 31 cells (control, MS1 single), n = 30 cells (BAPTA, MS1 single), n = 70 groups of cells (control, MS1 multiple) and n = 58 groups of cells (BAPTA, MS1 multiple), n = 12 cells (control, A375), n = 9 cells (BAPTA, A375) * P = 0.0351, **** P < 0.0001)

To investigate whether Ca^2+^ elevation is sufficient to trigger RhoA oscillations and to explore the underlying mechanisms, we induced persistent Ca^2+^ increase in HUVECs, MS1 and A375 cells using a Ca^2+^ ionophore ionomycin, in the absence of physical confinement. We again observed oscillatory RhoA activity in these cell lines, occurring at the same average frequency (approximately 0.6 hours^-1^) as that observed during cell migration under physical confinement (**Figure 2A, Supplementary Figures 2A-C and Methods S2**). Moreover, the bulk, population-level immunoblotting experiments with HUVECs, MS1 and A375 cells also revealed oscillatory changes in the phosphorylation of a downstream RhoA effector, the phosphorylated Myosin Light Chain (pMLC), suggesting that Ca^2+^-triggered RhoA oscillations occur synchronously across the cell population over at least 3 cycles (**Figure 2B and Supplementary Figures 2D,E**). Interestingly, we also observed a rapid decrease in acetylation of α-tubulin, potentially indicating dynamically changing the stability of microtubules (MT) (Maruta et al., 1986). This result raised the possibility that the onset of the dynamic changes in the RhoA activity may be triggered by a reduction in MT stability. The RhoA regulator whose activity depends on MT stability is a RhoA Guanine nucleotide Exchange Factor (GEF), GEF-H1 (Krendel et al., 2002). This RhoA GEF protein is bound to MT in an inactive state, but is activated following dissociation from MTs, e.g., during MT depolymerization. We indeed found that GEF-H1 activity oscillated in phase with the cycles of RhoA activity and MLC phosphorylation (**Figure 2B and Supplementary Figures 2D,E**). Surprisingly, GEF-H1 abundance also oscillated with the same frequency, but in the phase opposite to that of the RhoA oscillations. This result suggested that GEF-H1 could be a critical determinant of the oscillatory RhoA dynamics, and the resulting cell motility. We indeed observed that, in A375 cells, a shRNA-based knockdown of GEF-H1 expression resulted in abrogation of RhoA and pMLC oscillations (**Figure 2E**), reduction of Myosin-II mediated force generation (**Supplementary Figure 3**) (Munevar et al., 2001), and a severe reduction in the cell migration speed under physical confinement (**Figures 2C,D**). In combination, these results strongly implicated GEF-H1 in regulation of Ca^2+^-mediated oscillatory dynamics of RhoA and raised the question of how the activity and abundance of GEF-H1 are controlled in an oscillatory fashion.

**Figure 2.**
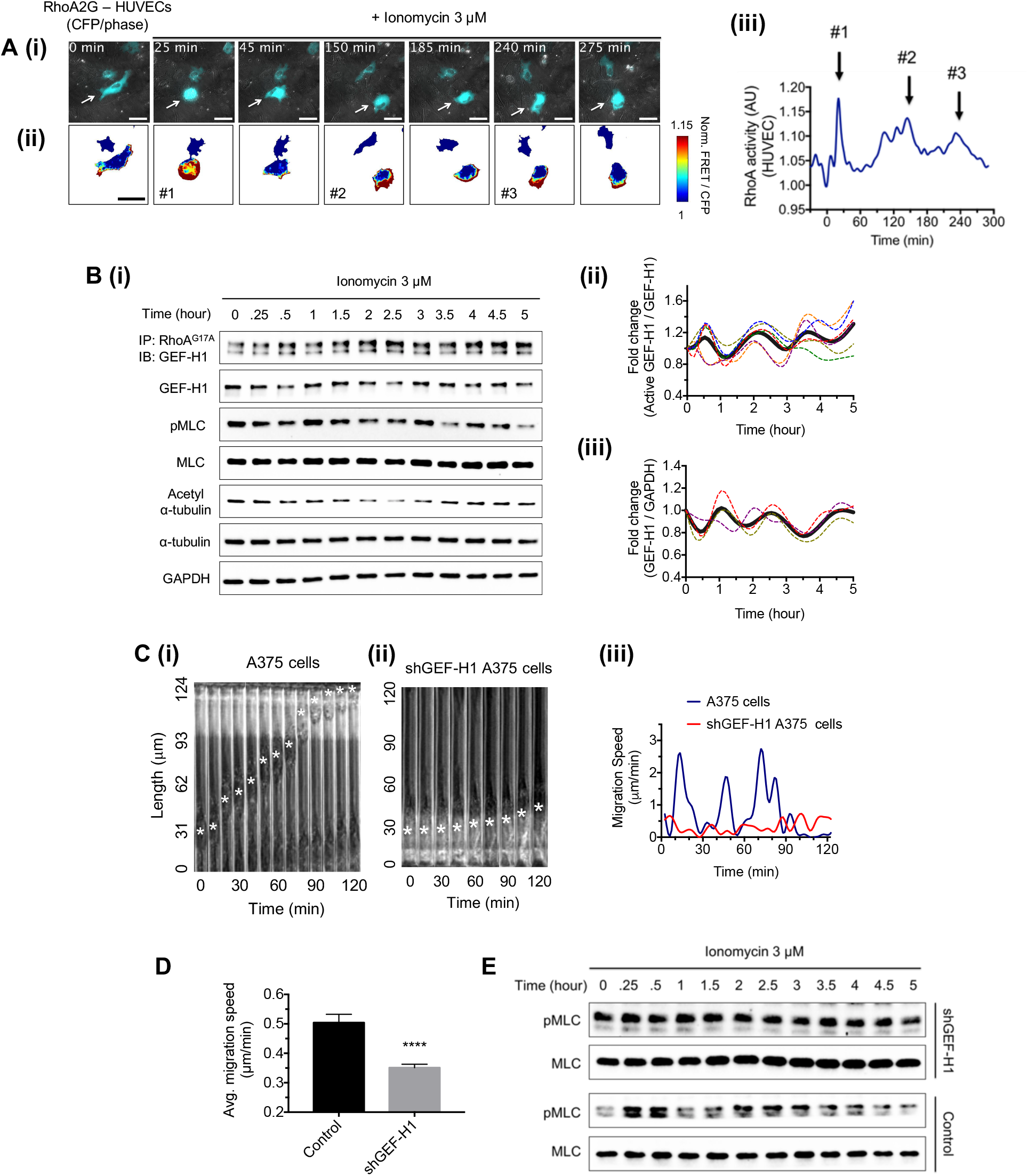
An increase of intracellular Ca^2+^ concentration by ionomycin promotes GEF-H1 dependent RhoA oscillations without spatial cell confinement. **(A)** (i) Representative time-lapse CFP images of RhoA2G – HUVECs superimposed on phase-contrast images and (ii) the corresponding pseudo-colored images of RhoA activity. The color bar indicates the ratio of the FRET signal over CFP. Scale bar: 50 μm. (iii) RhoA activity of the cell with the arrow in A (i). The numbers indicate peaks that were identified by the peak detection algorithm (Supplementary Figure 2A). **(B)** (i) Immunoblot of activation level and total abundance of GEF-H1, and of pMLC, MLC, acetylated a-tubulin and a-tubulin in HUVECs treated by ionomycin (3 μM). GAPDH was used as a loading control. (ii) Fold-change quantification of the active GEF-H1 level normalized by the total GEF-H1 abundance and (iii) total GEF-H1 level normalized to GAPDH. The dotted colored curves indicate results of independent experiments and the gray solid curves represent the averages of the dotted lines. The curve was created by the method of cubic spline interpolation using the function in Prism7. **(C)** A time-lapse kymograph tracking (i) A375 cell and (ii) shGEF-H1 A375 cell migrating in a spatially confining channel (version1). Asterisks indicate the nuclei of the cells. (iii) Migration speed dynamics of cells shown in (i) and (ii). **(D)** Average migration speed for shGEF-H1 A375 cells (n = 497) contrasted with that of un-infected A375 cells (n = 243). Data were analyzed by unpaired two-tailed t-test with error bar representing s.e.m. (**** P < 0.0001) **(E)** Immunoblot of pMLC and MLC in shGEF-H1 A375 and un-infected A375 cells in response to treatment with ionomycin (3 μM).

### Molecular mechanisms controlling dynamic alterations of the molecular clock components

Our results suggest that an increase in intracellular Ca^2+^ can initially trigger elevated GEF-H1 activity through enhanced MT depolymerization. This hypothesis is consistent with prior publications (Weisenberg, 1972; Weisenberg and Deery, 1981), suggesting that elevation of intracellular Ca^2+^ can reduce MT stability. To explore this further we directly assessed MT polymerization by tracking a commonly used plus-end tracking protein, EB1 tagged with GFP in HUVECs. To facilitate the analysis, the cells were examined on a nano-structured substratum (Kim et al., 2012; Park et al., 2019), previously shown to elicit anisotropic MT polymerization and cell morphology (Tabdanov et al., 2018). Tracking EB1 dynamics (see **Methods** for details) yielded a number of MT polymerization characteristics, revealing in particular that Ca^2+^ depletion with BAPTA resulted in an increased length of persistent growth of MT polymers, whereas Ca^2+^ upregulation with ionomycin or thapsigargin (an inhibitor of the sarco/endoplasmic reticulum Ca^2+^ ATPase (SERCA) pumps) decreased this length (**Figures 3A,B**). Since, the average growth rate of MTs was not altered under any of these conditions (**Table S1**), we concluded that persistently elevated Ca^2+^ levels can indeed promote MT depolymerization. This conclusion was further supported by the observation that the axial ratios of the cellular shapes increased following BAPTA pretreatment but decreased following ionomycin and thapsigargin pretreatments (**Figure 3C and Supplementary Figure 4B**), consistent with the expectations based on the degree of MT polymerization (cf. the effect of colchicine in **Figure 3C**). Overall, the results argue that elevated Ca^2+^ levels can indeed result in MT depolymerization, and as a consequence, a transient increase in GEF-H1 activity.

**Figure 3.**
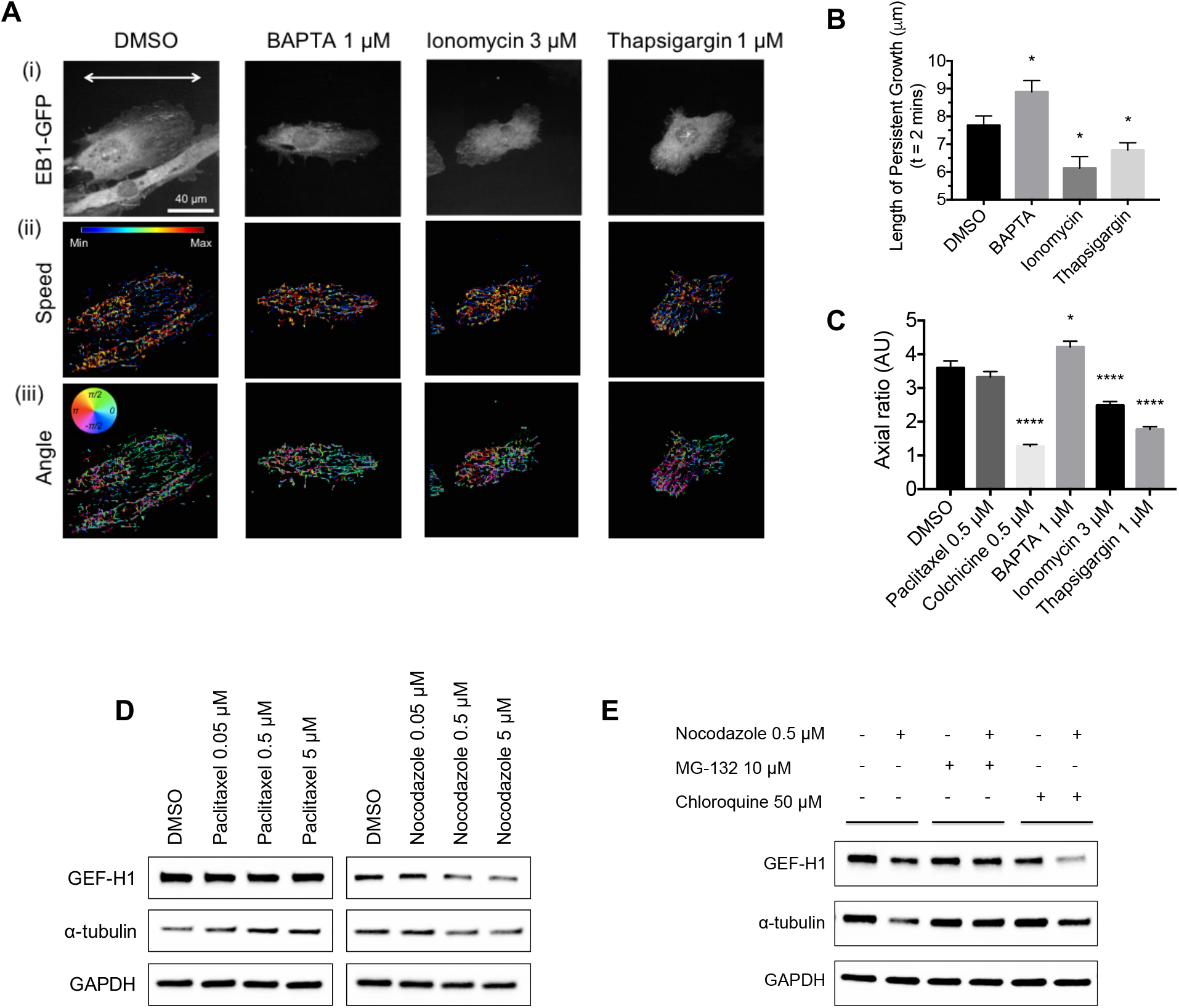
A persistent increase in intracellular Ca^2+^ leads to MT depolymerization, causing proteosomal GEF-H1 degradation. **(A)** (i) Representative fluorescent images of EB1-GFP-expressing HUVECs cultured on a nano-grooved patterned substratum following treatment by DMSO, BAPTA (1 μM), ionomycin (3 μM) and thapsigargin (1 μM) for 15 hours. The white arrow indicates the direction of the grooves and ridges in the pattern. Scale bar: 40 μm. Results of an automated tracking analysis representing (ii) speed and (iii) angle of trajectories of EB1-GFP clusters from images taken every 2 seconds for 2 minutes. **(B)** Comparison of the length of persistent MT growth over 2 minutes for the conditions of treatment by DMSO, BAPTA (1 μM) (* P = 0.0401), ionomycin (3 μM) (* P = 0.0113) and thapsigargin (1 μM) (* P = 0.0447). Data were analyzed by unpaired two-tailed t test with error bar representing s.e.m. **(C)** Comparison of axial ratios of HUVECs under the conditions of treatment with DMSO (n = 106 cells), paclitaxel (0.5 μM) (n = 140 cells), colchicine (0.5 μM) (n = 132 cells), BAPTA (1 μM) (n = 135 cells), ionomycin (3 μM) (n = 130 cells) and thapsigargin (1 μM) (n = 113 cells). Data were analyzed by one-way ANOVA with error bar representing s.e.m. (* P = 0.0116, **** P < 0.0001). **(D)** Immunoblot of GEF-H1 and a-tubulin abundance in HUVECs following treatment with paclitaxel (0.05, 0.5, 5 μM) and nocodazole (0.05, 0.5, 5 μM) for 15 hours. GAPDH was used as a loading control. Analysis is based on 3 independent experiments. **(E)** Immunoblot of GEF-H1 and a-tubulin abundance in HUVECs following treatment with MG-132 (10 μM), chloroquine (50 μM) and nocodazole (0.5 μM) for 15 hours. GAPDH was used as a loading control. Analysis is based on 3 independent experiments.

We next explored why a persistent increase in intracellular Ca^2+^ levels can trigger oscillations in the GEF-H1 abundance, including the initial decrease following ionomycin treatment (**Figure 2B and Supplementary Figures 2D,E**). This effect may reflect a protective role of GEF-H1-MT binding, which may increase the stability of GEF-H1 by protecting it from degradation. Camdependent MT dissociation would thus expose GEF-H1 to enhanced degradation, acutely decreasing its abundance. Indeed, we found that cell pre-treatment with MT-destabilizing agents, nocodazole and colchicine decreased GEF-H1 abundance in a dose and time dependent manner (**Figure 3D and Supplementary Figures 4C-E**). We then re-examined the effects of these MT destabilizing agents in the presence of inhibitors of proteasomal (MG-132) or lysosomal (Chloroquine) degradation. We found that inhibition of proteasomal degradation stabilized GEF-H1 abundance in the presence of both nocodazole and colchicine (**Figure 3E and Supplementary Figure 4F**). On the other hand, inhibition of lysosomal degradation did not prevent GEF-H1 degradation, and slightly enhanced the effects of both MT destabilizing agents. Furthermore, inhibiting proteasomal degradation in a high intracellular Ca^2+^ condition abrogated oscillations for both GEF-H1 and pMLC (**Supplementary Figure 4G**), indicating that RhoA oscillation is enabled by proteasome-mediated GEF-H1 degradation driven by MT depolymerization.

GEF-H1 abundance is restored in the second phase of oscillation. This effect could result from augmentation of GEF-H1 synthesis that may be dependent on persistent Ca^2+^ increase. To test this hypothesis, we pharmacologically modulated the intracellular Ca^2+^ by treatment with BAPTA, ionomycin or thapsigargin over a prolonged period of time. We found that a long term (15 hours) Ca^2+^ elevation by both ionomycin and thapsigargin led to an increase in the abundance of GEF-H1 to the levels beyond the maximal levels observed during the shorter-term (5 hours) oscillatory responses in HUVECs, MS1 and A375 cells (**Figure 4A, Figure 5D and Supplementary Figure 5B,C**). We further found an approximately 2.5-fold increase in *GEF-H1* mRNA levels following 15 hours’ cell treatment by thapsigargin (**Figure 4E**). This result suggested that elevated intracellular Ca^2+^ up-regulated GEF-H1 through increasing *GEF-H1* transcription rate. There are multiple transcriptional factors whose activity is modulated by Ca^2+^ inputs, including NFAT, MEF2 and CREB (Dolmetsch et al., 1998; Mao et al., 1999; Sheng et al., 1991; Youn et al., 1999). We therefore tested if the transcriptional *GEF-H1* regulation could be attenuated by inhibitors of these pathways (Fish et al., 2017; Kolovos et al., 2016; Noren et al., 2016; Sacilotto et al., 2016). We found that neither VIVIT, a specific and selective inhibitor of the NFAT signaling pathway, nor MEF2 activity inhibitor BML-210 had any detectable effect on GEF-H1 abundance (**Figures 4C,D**). On the other hand, selective inhibition of CREB regulation by the compound 666-15 partially reversed upregulation of *GEF-H1* expression and GEF-H1 abundance in response to Ca^2+^ elevation by thapsigargin (**Figure 4B,E**). These experiments suggested that GEF-H1 abundance can be upregulated through an increased synthesis in a Camdependent fashion. Overall, these results paint a complex picture of Ca^2+^-dependent control of GEF-H1 expression and activity (**Figure 4F**). On the one hand, elevated Ca^2+^ levels can reduce MT polymerization and thus acutely lead to an increase in the GEF-H1 activity, while also targeting it for proteasomal degradation. On the other hand, over a longer term Ca^2+^ input can upregulate GEF-H1 synthesis, which can lead to a transient increase in GEF-H1 abundance in the second part of each oscillation cycle.

**Figure 4.**
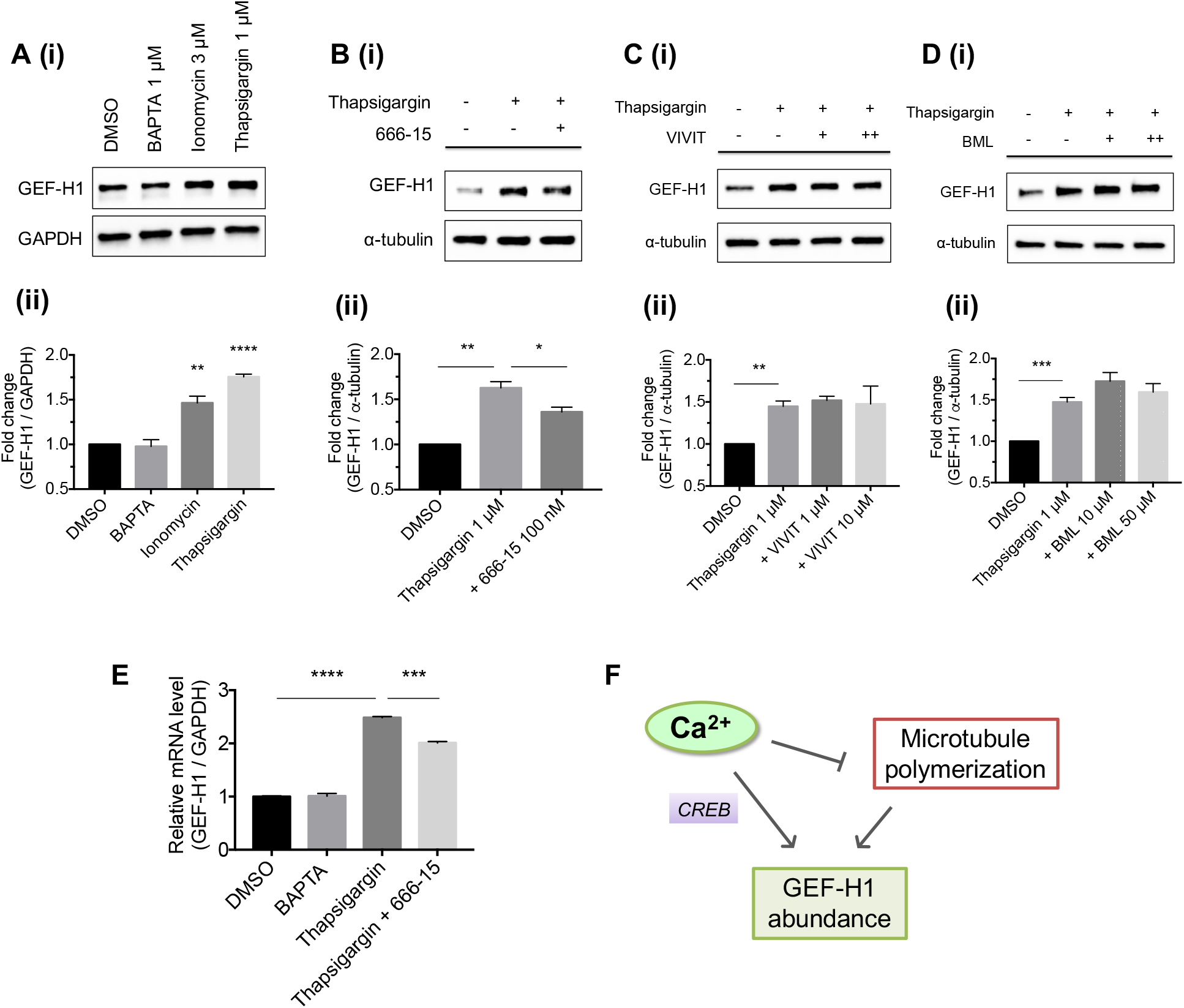
Persistently elevated intracellular Ca^2+^ enhances CREB-dependent GEF-H1 transcription. **(A)** (i) Immunoblot of GEF-H1 abundance in HUVECs following treatment with DMSO, BAPTA (1 μM), ionomycin (3 μM) and thapsigardin (1 μM) for 15 hours. GAPDH was used as a loading control. Analysis is based on 3 independent experiments. (ii) Fold-change quantification of GEF-H1 abundance normalized over GAPDH (** P = 0.0034, **** P < 0.0001). Data were analyzed by unpaired two-tailed t-test with the error bar representing s.e.m. **(B-D)** (i) Immunoblot analysis (ii) and fold-change quantification of GEF-H1 in HUVECs following treatment of DMSO, thapsigargin (1 μM) and/or (B) 666-15 (100 nM) (** P = 0.005, * P = 0.0208), (C) VIVIT (1, 10 μM) (** P = 0.0021) and (D) BML (10, 50 μM) (*** P = 0.0001) for 15 hours. a-tubulin was used as a loading control. Data were analyzed by unpaired two-tailed t test with error bar representing s.e.m. Analysis is based on 3 independent experiments. **(E)** GEF-H1 mRNA expression level change vs. GAPDH in HUVECs following treatment with BAPTA (1 μM), thapsigargin (1 μM) and/or 666-15 (100 nM) for 15 hours. Data were analyzed by unpaired two-tailed t test with error bar representing s.e.m. (**** P < 0.0001, *** P = 0.0001) **(F)** Schematic diagram summarizing the effect of intracellular Ca^2+^ on microtubules and GEF-H1 abundance.

### Negative feedback regulation mechanisms in the molecular clock regulation

A key feature of most biological oscillations is the presence of negative feedback regulation. We therefore explored putative mechanisms of such feedback loops in the control of GEF-H1 abundance and activity, and the resulting oscillatory RhoA dynamics. Our experimental results suggested both GEF-H1 activity and abundance can be controlled by association with MTs, and thus MT stability (Krendel et al., 2002). The feedback loop can be closed if GEF-H1 can, in turn, have an effect on MT polymerization. To explore this possibility, we examined the EB1 dynamics following GEF-H1 knockdown (**Figure 5A**). Imaging EB1 in both GEF-H1 knockdown cells and cells expressing a scrambled shRNA control revealed no difference in average MT growth rate across WT and GEF-H1 knockdown cells (**Table S2**). However, GEF-H1 knockdown resulted in a longer length of persistent MT growth (**Figure 5B**), indicating that GEF-H1 can promote MT depolymerization, indeed constituting a putative negative feedback loop.

**Figure 5.**
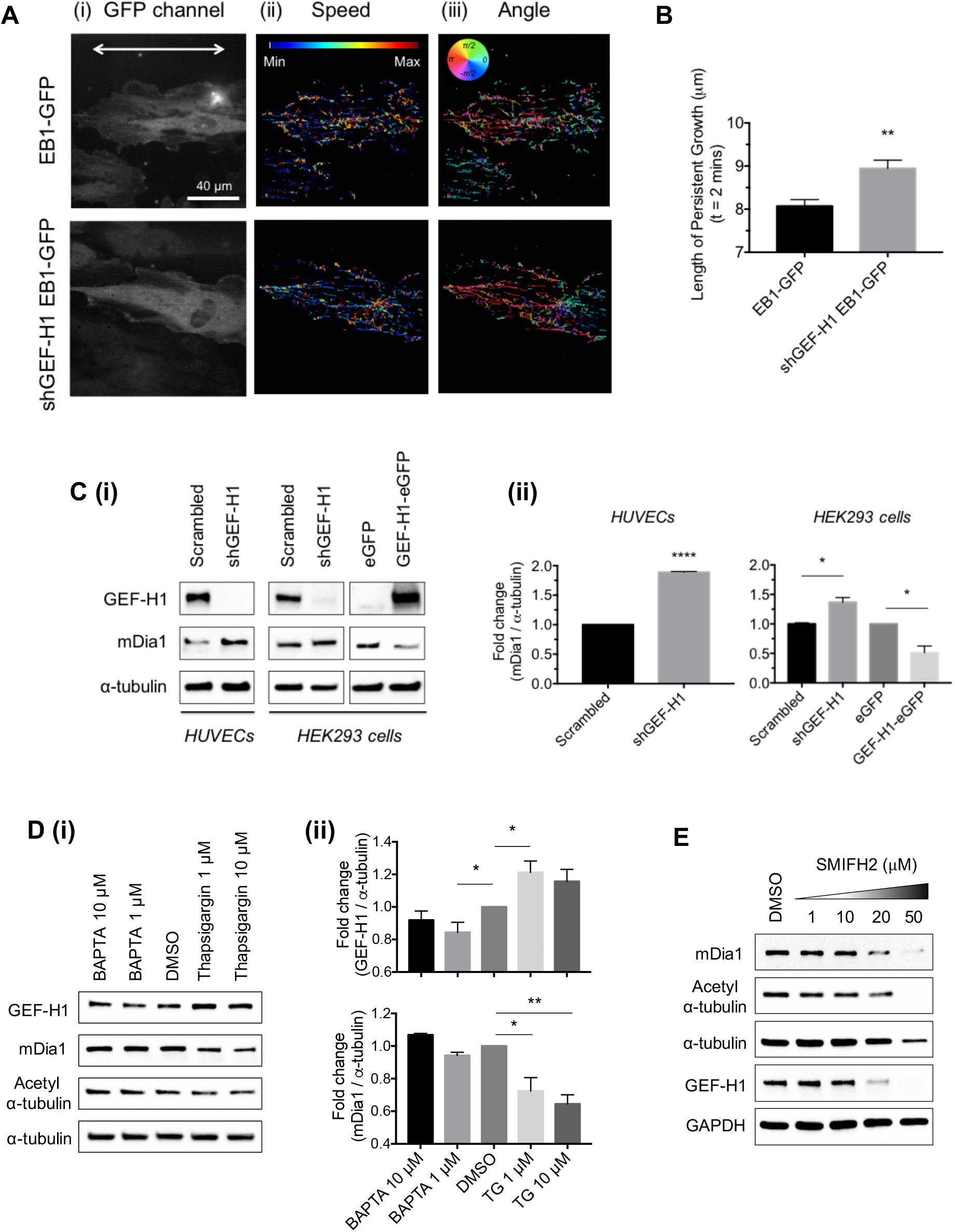
GEF-H1 abundance negatively regulates mDia1-mediated microtubule polymerization. **(A)** Representative fluorescent images of EB1-GFP and shGEF-H1 EB1-GFP expressing HUVECs on a nanogrooved patterned substratum. The white arrow indicates the direction of the grooves and ridges in the pattern, analyzed as in Fig. 3. Scale bar: 40 μm. Results of an automated tracking analysis representing (ii) speed and (iii) angle of trajectories of EB1-GFP clusters from images taken every 2 seconds for 2 minutes. **(B)** Comparison of length of persistent growth of microtubules over 2 minutes determined using EB1-GFP in un-infected (n = 13 cells) and shGEF-H1 expressing (n = 23 cells) HUVECs. Data were analyzed by unpaired two-tailed t test with error bar representing s.e.m. (** P = 0.0036) **(C)** (i) Immunoblot analysis of GEF-H1 and mDia1 abundances in cells infected with scrambled shRNA, shGEF- Hl (HUVECs, HEK293 cells) or a construct encoding the GEF-H1-eGFP (HEK293 cells). a-tubulin was used as a loading control. Analysis is based on 3 independent experiments. (ii) Fold-change quantification of mDia1 expression relative to a-tubulin. Data were analyzed by unpaired two-tailed t test with error bar representing s.e.m. (shGEF-H1 HUVECs **** P < 0.0001, shGEF-H1 HEK293 cells * P = 0.0460, GEF-H1- eGFP HEK293 cells * P = 0.0499) **(D)** (i) Immunoblot analysis of abundances of GEF-H1, mDia1, acetylated a-tubulin and a-tubulin in MS1 cells following treatments with DMSO, thapsigargin (1, 10 μM) and BAPTA (1, 10 μM) for 15 hours. (ii) Fold-change quantification of expression relative to a-tubulin of GEF- H1 (BAPTA 1 μM: * P = 0.0353, thapsigargin 1 μM: * P = 0.0167) and mDia1 (thapsigargin 1 μM: * P = 0.028, thapsigargin 10 μM: ** P = 0.0031) for the experiments shown in (i). Images are representative of 3 independent experiments. **(E)** Immunoblot analysis of the abundances of mDia1, acetylated a-tubulin, a-tubulin and GEF-H1 in HUVECs following treatment with DMSO and SMIFH2 (1, 10, 20, 50 μM) for 15 hours. GAPDH was used as a loading control. Analysis is based on 3 independent experiments.

To further address the mechanisms for GEF-H1-mediated increase in MT depolymerization, we focused on a RhoA (and thus GEF-H1) effector, formin mDia1, previously shown to regulate both actin polymerization and MT stability (Bartolini et al., 2016; Palazzo et al., 2001). We found that increasing doses of pharmacological inactivation of mDia1/2 by a specific inhibitor, SMIFH2, led to a progressively decreasing a-tubulin acetylation and GEF-H1 abundance (**Figure 5E**). This result thus indeed suggested that mDia1/2 can mediate a negative feedback controlling GEF-H1 abundance. We then explored whether the mDia1 abundance may also be controlled by GEF-H1. We found that GEF-H1 knockdown in HUVECs and HEK293 cells, the expression level of mDia1 was higher vs. cells expressing the scrambled shRNA control cells (**Figure 5C**). Conversely, overexpression of GEF-H1 in HEK293 cells resulted in a lower expression level of mDia1 (**Figure 5C**). Furthermore, we found that cell treatments with thapsigargin or ionomycin led to both an increase of GEF-H1, and a concurrent decrease in mDia1 levels (**Figure 5D and Supplementary Figures 5B**). These data indicated that elevated GEF-H1 expression can have a negative effect on the mDia1 abundance. This effect was in apparent contradiction with the expected positive effect of GEF-H1 activity on the activity of mDia1, through upregulation of the activity of RhoA (Watanabe and Higashida, 2004). This contradiction is however reconciled by our earlier observation that GEF-H1 activity and abundance oscillate in opposite phases, and are therefore anti-correlated. Therefore, the positive effect of GEF-H1 activity on mDia1 activity, and the negative effect of GEF-H1 abundance levels on mDia1 concentration can be resolved in time in different phases of oscillations.

### The dual feedback mechanism can explain the oscillatory dynamic responses of the molecular clock

Our analysis paints a complex picture of the regulatory network putatively controlling GEF-H1 activity and abundance, and RhoA activity oscillations, in a Ca^2+^ dependent fashion. In particular, our data reveal two coupled regulatory negative feedbacks controlling these oscillations in a MT and mDia1 dependent fashion (**Figure 6A**). These feedback loops are both regulated in a MT-dependent fashion, but the first loop accounts for regulation of GEF-H1 expression, positively controlled by MT stabilization, and the second – for regulation of GEF-H1 activity, negatively controlled by MT stabilization. The first loop reflects the negative effect of an increased GEF-H1 abundance on the abundance of mDia1, whereas the second loop recapitulates the positive effect of GEF-H1 activity on the RhoA-dependent activity of mDia1 (Watanabe and Higashida, 2004). mDia1 activity and expression levels can both positively affect MT stability thus closing both feedback loops and coupling them into the same circuit. This molecular circuit can be stimulated and controlled by elevated Ca^2+^ through its effects on both MT stability and CREB-dependent GEF-H1 transcriptional up-regulation. This postulated mechanism can be validated if it can faithfully reproduce not only the observed dynamics of GEF-H1 and RhoA oscillations, but also the effects of further perturbations of the molecular circuit. To explore this, we first expressed the above mechanism in terms of a mathematical model (**Figure M1 in Methods S3**). We found that this model can indeed display oscillatory dynamics, and account for the opposite phases of GEF-H1 activity and abundance (**Figure 6B** and **Figure M2 in Methods S3**). We further ‘trained’ the model using the dynamical data observed above, ensuring that the model can display the observed average oscillation frequency and phases for the diverse circuit components. We then tested the proposed mechanism expressed as this mathematical model by examining its ability to predict the behavior of other model components and effects of additional perturbations of the molecular circuit identified here.

**Figure 6.**
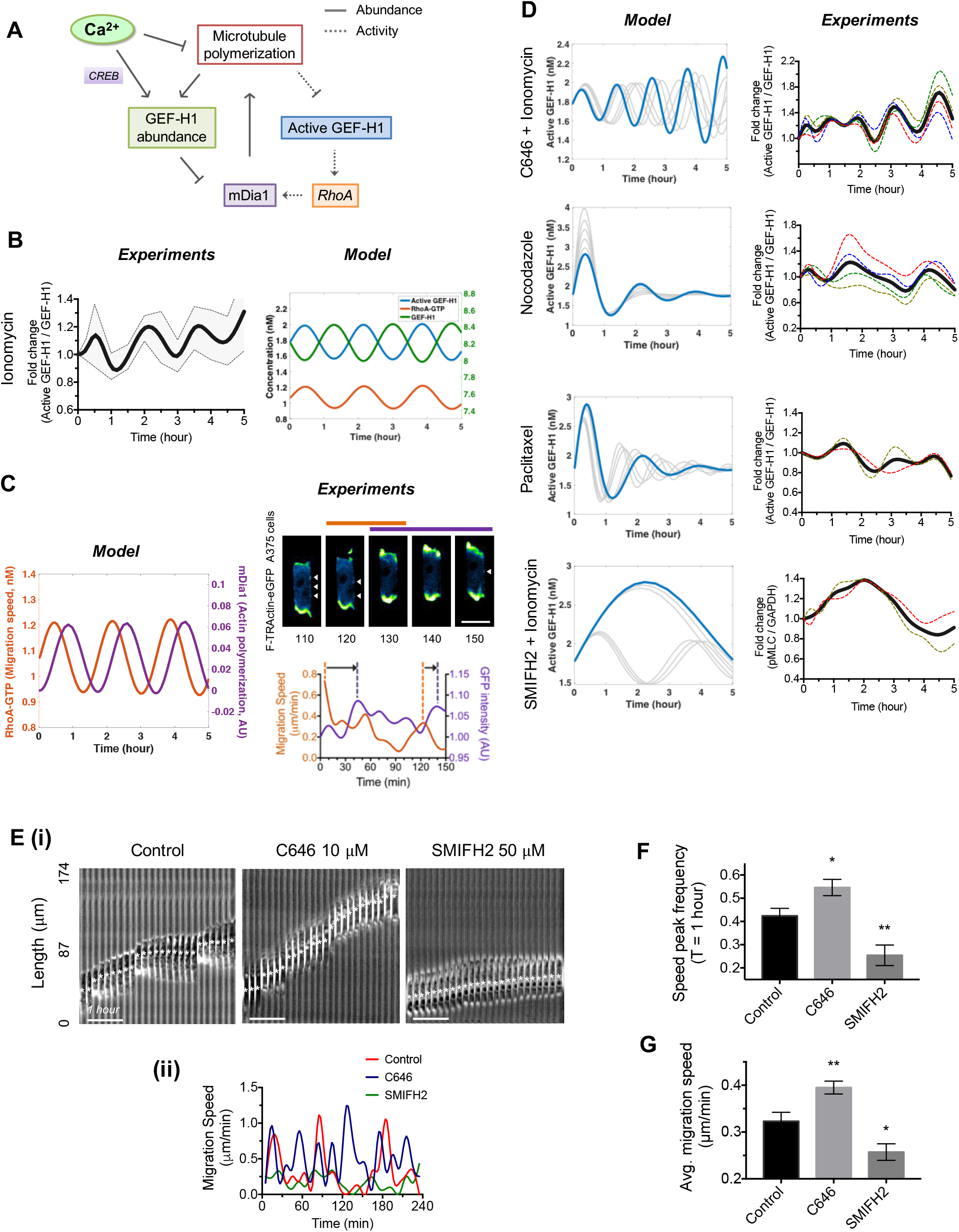
The mechanism underlying the molecular clock driving cell migration under spatial confinement. **(A)** Schematic diagram of the putative molecular network driving the molecular clock. **(B)** Experimental results (left) of the temporal dynamics of GEF-H1 activity in response to treatment with ionomycin (from Figure 2B (ii)) and model prediction (right) of temporal dynamics of the GEF-H1 activity (blue, left y-axis), RhoA-GTP abundance (red, left y-axis) and GEF-H1 abundance (green, right y-axis) **(C)** Model prediction (left) and experimental results (right) of a time delay between oscillatory RhoA activation (assayed as the migration speed) and mDia1 activity (assayed as actin polymerization). Time lapse images of F-TRActin-eGFP A375 cells migrating under confinements in narrow channels (version1) (top right). Horizontal bars indicate the timing of the migration speed (orange) and GFP intensity (purple) peaks corresponding to a portion of the graph below (bottom right). White arrows indicate blebs at the lateral cell periphery. Scale bar: 10 μm. Quantification of the migration speed and the intensity of actin filamentation over time for this experiment is shown in bottom right. **(D)** The effects of pharmacological perturbations of the molecular clock analyzed by mathematical model prediction (left) and experimental validation (right). Gray lines indicate simulation results for various model parameters (left). The dotted color lines indicate independent experimental results and the dark gray line represents averages of these experimental data (right). **(E)** (i) Time-lapse kymograph tracking A375 cells under confinements (version1) following treatments with C646 (10 μM) and SMIFH2 (50 μM) for 15 hours. Scale bar: 1 hour. (ii) Migration speed dynamics of the cells in (i). **(F)** Migration speed peak frequency over 1-hour (*P = 0.0202, **P = 0.0064) **(G)** Average migration speed for A375 cells treated as in (E) and untreated control cells (*P = 0.0214, **P = 0.0067). Data were analyzed by unpaired two-tailed t test with error bar representing s.e.m. (n=10 each)

As a first test and to further characterize the relevance of the proposed mechanisms to cell migration, we explored the predicted dynamic behavior of mDia1 activity. The model suggested that mDia1 displays oscillatory activation with a phase shift of approximately 30 minutes vs. the oscillatory activity of RhoA (**Figure 6C**). We then explored the dynamic mDia1 activity taking advantage of its formin function, promoting actin polymerization. To enable this analysis, we probed filamentous actin in A375 cells using F-TRActin conjugated with eGFP (Watanabe and Higashida, 2004). We found that A375 cells, under physical confinement, displayed actin-rich bleb formation, mostly at the front and rear, but also, in multiple smaller blebs, on the sides of the cell bodies, in apparent tight contact with the channel walls (**Figure 6C and Supplementary Figure 6A,B**). This actin localization pattern was in contrast to preferential actin localization to stress fibers in the 2D migration controls (**Supplementary Figure 6A**), and suggested the so-called ‘chimneying’ mechanism of cell migration (Paluch et al., 2016). We then measured both the dynamic changes in the migration speed and the intensity of actin filamentation during cell migration. During cell migration, the small blebs at the sides of the cells tended to remain in the same positions as cells underwent forward translocation phases, again suggesting that these blebs were points of force application in the ‘chimneying’ migration (**Figure 6C and Supplementary Figure 6B**). Importantly, we found that actin filamentation was pulsatile, with the peaks of F-actin abundance in the blebs following the peaks of cell migration speed (indicative of peak RhoA activity values) with the delay of approximately 30 minutes. This result was consistent with the model prediction under the assumption that actin filamentation in this migration mode is driven by the formin activity associated with F-actin rich bleb formation.

### The frequency of the molecular clock can be modulated in a predictable manner controlling the speed of cell migration

To further validate the proposed molecular clock through experimental perturbations of the underlying biochemical components, we first analyzed the mathematical model to determine the sensitivity of the observed oscillatory behavior to variation of the model parameters (**Figure M3 in Methods S3**). We found that the frequency and amplitude of oscillations were predicted to be most sensitive to the perturbations of MT stability and mDia1 expression/activity levels. These parameters are indeed central to both negative feedback loops postulated in the proposed mechanism. To validate this prediction, we therefore focused on perturbations of MT stability and of mDia1 activity. Our data so far suggested that MT stability is correlated and may be modulated its acetylation in the presence of elevated Ca^2+^ levels. Although the role of MT acetylation in controlling MT stability is still a subject of intense investigation, we indeed found that MT acetylation levels were correlated with the MT stability under various experimental conditions (**Supplementary Figures 5B,C and Supplementary Figures 7A-D**). This result raised the possibility that control of MT acetylation may provide a way to perturb MT stability, which may be complementary to the use of more common drugs, such as colchicine and paclitaxel. To further test the putative causal role of MT acetylation in determining MT stability, we perturbed the expression of aTAT-1/MEC-17, the key MT acetyltransferase, through inhibition of a negative regulator of its expression, another acetyltransferase p300 (Mackeh et al., 2014). We found that inhibition of p300 activity using C646, indeed upregulated both aTAT1/MEC-17 expression and MT acetylation (**Supplementary Figures 7F**). We then examined MT stability following the same p300 perturbation, finding it to be significantly increased (**Supplementary Figures 7G,H**), the result that strongly suggested that elevated MT acetylation can indeed promote MT stability in the cells analyzed here. This result was further supported by the observation that the decrease of MT acetylation and stability in response to thapsigargin was rescued if p300 was also inhibited (**Supplementary Figure 7E**). Furthermore, since p300 is also involved in regulation of CREB mediated transcription (through modulation of the binding of the CREB-binding protein, CBP) (Bowers et al., 2010), we also examined GEF-H1 abundance for different doses of C646. We indeed found that the thapsigargin-mediated increase in GEF-H1 abundance was indeed reduced in a dose-dependent fashion (**Supplementary Figure 7E**). This result underscored the dual effect of p300 in controlling the acetylation-mediated biochemical processes modulating both MT stability and GEF-H1 synthesis.

We then examined the mathematical model for predictions of the effects of perturbations of the MT stability. The model predicted that an increase in MT stability and decrease of GEF-H1 synthesis (modeling the effect of inhibition of p300) in the presence of elevated Ca^2+^ inputs would lead to a pronounced increase in the oscillation frequency and a gradual rise in the oscillation amplitude vs. the effect of elevated Ca^2+^ levels alone (**Figure 6D**). We indeed found both these effects in the experimental analysis of the ionomycin-triggered Ca^2+^ oscillations observed in the presence of the p300 inhibition (**Figure 6D and Supplementary Figure 8A**). We then examined if the oscillations could be directly triggered by MT modifying agents, without Ca^2+^ elevation. The model predicted that both exogenous MT stabilization and destabilization could each trigger transient oscillatory responses, albeit with oscillations being damped (amplitude gradually decreasing in time) (**Figure 6D**). We indeed found that cell treatment with nocodazole or paclitaxel could both trigger oscillations, although the experimental degree of damping was lower than that predicted by the model (**Figure 6D and Supplementary Figure 8B,C**). These results provided support for both the proposed mechanism and, more specifically, the key role of MT stability in controlling the oscillation dynamics. Finally, the model predicted that inhibition of mDia1 would lead to a pronounced decrease in the oscillation frequency (**Figure 6D**). We indeed found that pretreatment of cells with SMIFH2 led to a much slower oscillatory response (**Figure 6D and Supplementary Figure 8D**). In combination, our analysis suggested that the model could correctly predict the dynamic effects of various perturbations of the regulatory network postulated here as the key mechanism. These findings further supported the proposed mechanism underlying the oscillatory GEF-H1 and RhoA dynamics. With these results in hand, we then explored the functional significance of the observed oscillations in controlling cell migration in physically confining environments.

Our results suggest that elevation of intracellular Ca^2+^, occurring as a consequence of cell movement in physically confining 3D environments, can trigger a molecular clock regulating oscillations in RhoA activity. Furthermore, our integrative mathematical and experimental analysis predicted that the oscillation frequency can be strongly modulated by perturbations of MT stability and mDia1 activity. To test if these changes in the clock activity would translate into the corresponding alterations of cell migration dynamics, we investigated cell migration in physically confining environments following cell pre-treatment with p300 and mDia1 inhibitors (**Figure 6E**). Strikingly, we found that an increase in RhoA oscillation frequency in response to p300 inhibition led to correspondingly more frequent oscillatory changes in the cell migration speed and a pronounced enhancement of the average migration speed and the distance travelled by the cells vs. the untreated control (**Figures 6E-G**). The inhibition of mDia1 had the opposite effect, leading to a significantly slower cell migration vs. the control (**Figures 6E-G**). Overall, these results strongly suggested that the observed oscillations in RhoA activity can have a profound effect on key properties of cell migration in confined spaces, fully consistent with the mechanism postulating two coupled GEF-H1 dependent negative feedbacks triggered and maintained by elevated intracellular Ca^2+^ levels.

## Discussion

The biomolecular mechanisms controlling cell migration are complex and only partially understood. Our analysis suggests a mechanistic view of how microtubule and actin cytoskeleton rearrangements, required for efficient cell locomotion in confined spaces, are dynamically controlled in a cyclic fashion. In particular, according to the putative mechanism of 3D cell migration proposed here, mDia1-regulated actin and microtubule polymerization alternate with microtubule depolymerization and myosin-mediated actin cytoskeleton contraction, propelling cell movement. The key molecular component controlling this dynamical behavior is a RhoA GEF protein, GEF-H1, whose activity and abundance are regulated through two coupled negative feedbacks, both of which depend on association of this molecule with microtubules. The feedback loops are synchronized, determining the pace of the coordinated cycles of RhoA activity and microtubule polymerization-depolymerization. The biochemical interactions in this feedback-rich circuit can be strongly enhanced by a persistent increase in intracellular Ca^2+^ levels, triggered either due to mechanical inputs during the 3D migration through dense ECM matrices, or pharmacological means or, potentially, other biochemical inputs, such growth factor ligands. The Ca^2+^-mediated inputs regulate both the microtubule stability and elevated synthesis of GEF-H1, thus impinging on the regulatory network in two distinct ways. The cyclic activity of the circuit components can also be triggered through other circuit perturbations, including, e.g., induced microtubule depolymerization.

The molecular circuit analyzed in this study provides yet another example of the importance of rhythmic activity in biological systems (Cheong and Levchenko, 2010). In studying the mechanisms underlying these biological oscillators, it is important to demonstrate perturbations that not only abrogate oscillatory activity itself, but also modulate the frequency of activity cycles, and importance of this frequency for controlling key associated biological phenotypes. Predictive mathematical models can help perform this analysis by zeroing on the points of sensitivity and control of the underlying molecular mechanisms. In the analysis presented here, perhaps the most striking observation was the ability of the mechanistic model to predict the perturbations that had very strong effects on both the frequency of the molecular clock and the associated phenotype of cell migration speed. This quantitative analysis thus presents a sophisticated dynamic test of the proposed mechanisms and also provides a tool for their detailed exploration and use. It also raises questions for further exploration, e.g., why the melanoma cells analyzed here do not move with the maximal possible speed they are capable of, and under what circumstances this speed can be dramatically increased or increased. Furthermore, as described below, this analysis elucidates the importance of specific molecular components engaged in the dynamic control of the molecular clock activity.

The regulated sequence of precisely timed events inherent in the cyclic activity of the molecular clock described here can ensure that various biomolecular processes occur in specific temporal phases, thus separating in time what may be incompatible regulatory events. For instance, the temporal coordination can ensure that the RhoA-mediated, actomyosin contraction-dependent motive force and the increase in the hydrostatic pressure needed for formation of blebs essential for 3D migration do not conflict with potentially opposing forces stemming from microtubule and actin polymerization. In particular, the proposed mechanism predicts that the mDia1 activity and thus actin polymerization occurs with a delay (of approximately one quarter of an oscillation cycle) with respect to the peak of the RhoA activity. The delayed actin polymerization initiation can then assist in expansion and stabilization of the blebs due to actin polymerization within these membrane structures and the resulting application of forces required for cell attachment to the surrounding substratum through the so-called ‘chimneying’. Indeed, we observed that actin polymerization showed a dynamic increase, peaking with a delay with respect to the peak of RhoA-associated motive force, and leading to formation of actin-filled blebs not only at the front of the cell, but also at the lateral cell periphery, where the blebs persisted in contact with the substratum, as cells underwent locomotion, likely being points of force application.

The modeling and experimental results of this study underscore the critical importance of microtubule dynamics in controlling the oscillation frequency and amplitude. These findings are consistent with the previously published reports that microtubule instability, particularly in the uropod, is essential for 3D cell migration (Bouchet and Akhmanova, 2017; Hind et al., 2016). Our results further suggest that the increased microtubule dynamics is essential for cyclic activation of the RhoA- and mDia1-mediated processes and thus effective cell migration in dense extracellular matrices. These results are also interesting to contrast with prior observations that microtubule destabilization could trigger oscillations of cell polarity during cell migration in models of 1D cell migration: narrow flat strips of extracellular matrix (Zhang et al., 2014) or aligned parallel fibers mimicking the strands of extracellular matrix (Park et al., 2017). In all these settings the results suggest that the microtubule dynamics is not only directly interfaced with the regulatory networks controlling the cytoskeleton organization, but also is critical for their cyclic activation on diverse time scales.

The activation of the molecular clock described here by persistent Ca^2+^ elevation also underscores the key role of this pleiotropic second messenger in promoting cell migration. It is notable that, as we showed previously, in endothelial cells Ca^2+^ can also promote cell migration regulating MLCK (Noren et al., 2016; Yokota et al., 2015), thus converging on the myosin regulation by ROCK considered here. Therefore, in invasive endothelial cells (e.g., Tip cells during angiogenesis), persistent Ca^2+^ increase triggered by VEGF and mechanical stimuli can have multiple effects on the contractile apparatus controlling cell locomotion. Ca^2+^ can also be upregulated by other inputs, including non-canonical Wnt signaling (Endo et al., 2015; O’Connell et al., 2013) in melanoma cells increasing their invasive potential. More generally, intracellular Ca^2+^ might be key integrator of extracellular chemical and mechanical signals, regulating both the activity and expression of many of the critical regulatory components driving invasive cell spread.

Given the key importance of cell migration in regions of dense extracellular matrix, including invasive migration of metastatic cancer cells, our analysis can also serve as a stepping stone towards evaluation or reevaluation of potential clinical interventions, limiting angiogenesis or cancer cell dissemination. For instance, inhibition of p300 has been proposed as potential anti-cancer therapy (Lasko et al., 2017). However, our results suggest that application of p300 inhibitors might enhance cell dissemination and thus perhaps promote initiation of metastasis, suggesting a way to determine the range of applicability of these putative anti-cancer therapeutics. Furthermore, we note that the critical role of GEF-H1 in the molecular clock of cell migration in confined spaces suggests the need to evaluate this protein as a potentially interesting target for intervention. This role is particularly supported by the observation that a knockdown of GEF-H1 led to abrogation of the oscillatory RhoA activity and had a very strong negative effect on the spatially confined locomotion of an invasive melanoma cell line. These examples suggest that the analytic framework for the quantitative analysis presented here may indeed lead to important preclinical research directions.

Overall, our findings suggest a mechanistic framework that can help understand invasive 3D cell migration through dense extracellular matrices and other forms of cell migration, and thus provide a useful stepping stone for studying and controlling the associated processes in physiological and pathological settings.

## Materials and Methods

### Cell culture

A375 melanoma, Mile Sven 1 (MS1, mouse pancreatic endothelial) and HEK293 cell lines were purchased from ATCC (CRL-1619, CRL-2279 and CRL-1573 respectively) and maintained in DMEM high-glucose media (Corning, 10-013-CV) with heat inactivated FBS 10 % and PenicillinStreptomycin (Life technologies, 15070063) 1 %. Human umbilical vein endothelial cells (HUVECs) were purchased from Yale Vascular Biology and Therapeutics program and maintained in Medium 199 (Life technologies, 11150-059) with heat inactivated FBS (Life technologies, 16140-071) 20 %, Endothelial cell growth supplement 0.03 mg/ml (Sigma-aldrich, E2759), Heparin sodium salts (Sigma-aldrich, H3149-100KU) 0.05 mg/ml, HEPES (Life technologies, 15630106) 10 mM, GlutaMAX supplement (Life technologies, 35050061) 1X and Antibiotic-Antimycotic (Life technologies, 15240112) 1 %. 0.1 % gelatin was coated for 10 minutes at 37 ***°C*** incubator before seeding HUVECs on the culture dish. All cell lines were cultured in humidified 37 °C and 5 % CO_2_ incubator.

### Chemicals and antibodies

The following chemicals were used: Ionomycin (Cayman, 10004974), BAPTA (Invitrogen, B6769), Thapsigargin (Invitrogen, T7459), Paclitaxel (VWR, T104-0005), Colchicine (Sigma, C9754), Nocodazole (Sigma, M1404), C646 (Cayman, 328968-36-1), VIVIT (Cayman, 249537-73-3), BML 210 (Cayman, 537034-17-6), Fluo-4 AM (Introgen, F14201), SMIFH2 (Tocris, 4401), MG- 132 (Sellekchem, S2619) and Chloroquine (Sigma, C6628).

The following primary antibodies were used for immunoblotting and co-immunoprecipitation: GEF-H1 (Cell Signaling, 4076S), acetylated a-tubulin (Santa Cruz, sc-23950), a-tubulin (Sigma, T6074), a-tubulin (Santa Cruz, sc-5274), GAPDH (Santa Cruz, sc-20357), Phospho Myosin Light Chain 2 (Cell Signaling, 3674S), Myosin Light Chain 2 (Cell Signaling, 8505S), mDia1 (Cell Signaling, 5486S), rabbit IgG (Santa Cruz, sc-2027). The following secondary antibodies were used for immunoblotting: ECL rabbit HRP-linked IgG (GE Healthcare, NA934) and ECL mouse HRP-linked IgG (GE Healthcare, NA931).

### Device fabrication

The device was designed in LayoutEditor to generate spatially confining channels and AutoCAD 2012 for the fabrication of 3D collective cell migration device. We fabricated the molds using standard SU8 lithography techniques where the photoresist was exposed to patterned UV from the designed mask, etched and finally cleaned. To generate both channels and media outlets, we used the mask aligner (SUSS MJB4 Mask Aligner) and a standard spin coater for a two-step fabrication process. Then, the mask was cleaned and PDMS mixture (Sylgard 184) was poured into the mold and baked at 80 °C for 2 hours. The PDMS devices were removed and cut into appropriate sizes and punched with holes for inlets by 5 mm or 8 mm biopsy punch (Miltex). We designed the channels with the parameters (width: 3, 6, 10, 20 and 50 μm, height: 10 μm and length: 200 μm) and used two versions of widths for confining experiments: version1 (width: 3 μm) and version2 (width 6 μm). The device used for the analysis of cell migration in 3D extracellular matrix is described in the Supplementary Methods File1.

### Immunoblotting

We washed cells with PBS with MgCl2 1 mM, CaCl2 1 mM three times and lysed in RIPA buffer mixed with Halt proteases and phosphatase inhibitor cocktail (Thermo Scientific). Lysed cells were rotated for 10 minutes and spun down at 15,000 rpm by table top centrifuge for 10 minutes at 4 °C. Protein concentration from lysates were equalized and subsequently followed by a denaturation in 4x Laemmli buffer mixed with 2-mercaptoethanol by boiling at 75 °C for 15 minutes. Lysates were then separated by SDS-PAGE using 4-20 % Mini-PROTEAN TGX precast gels (Biorad) and transferred onto 0.2 μm nitrocellulose membrane using Tran-Blot Turbo (Biorad) set at 2.5V, 25 A for 10 minutes. Transferred membranes were incubated in blocking solution (3 % bovine serum albumin in TBST (TBS + 1 % Tween 20)) for 1 hour at room temperature on rotating shaker and incubated with primary antibody in blocking solution at 4 °C overnight on rotating shaker. Next day, membranes were washed three times with TBST for 15 minutes each and subsequently incubated with HRP conjugated secondary antibody in blocking solution for 1 hour at room temperature on rotating shaker. Samples were washed three times again with TBST for 15 minutes each and treated with Biorad Clarity Western ECL substrates to amplify signals. All membrane images were performed in ChemiDoc MP imaging system (Biorad). All immunoblot images were quantified using ImageJ.

### Active Rho GEF assay

Active RhoA guanine exchange factor (GEF) pull down assay has been described in detail elsewhere (Garcia-Mata et al., 2006; Guilluy et al., 2011). After transforming GST-RhoA^G17A^ plasmid in DH5-a competent cells, we induced protein expression by adding IPTG 100 μM by shaking the culture at room temperature for 20 hours. Then, bacteria cells were lysed with lysis buffer (20 mM HEPES, 150 mM NaCl, 5 mM MgCl2, 1 % (vol/vol) Triton X-100, 1 mM DTT and protease inhibitor cocktail (Sigma)), followed by a sonication (Qsonica) for 1 minute with pulses of 1 second using 3-mm tip set at 40 % and the lysate was clarified by centrifugation at 15,000 rpm for 15 minutes. The supernatant was transferred to a fresh tube and added with 500 μl of 50 % (vol/vol) GST tagged glutathione-sepharose slurry (GE Healthcare) in lysis buffer and rotated at 4 °C for 1 hour. After washing with lysis buffer and with HBS buffer (20 mM HEPES and 150 mM NaCl) twice each, the residual buffer was carefully aspirated and 0.5 volumes of glycerol was added with 0.02 % sodium azide.

For active Rho GEF pull-down assay, cells with 70 – 90 % confluency, a total of ~ 3.0 × 10^6^ cells, were lysed in Rho GEF buffer (20 mM HEPES, 150 mM NaCl, 5 mM MgCl2, 1 % (vol/vol) Triton X100, 1 mM DTT and protease inhibitor cocktail (Sigma)). After rotating cells for 10 minutes, cells were spun down with table top centrifuge set at 15,000 rpm for 10 minutes in a prechilled rotor. After equalizing protein concentration, 10 μl of GST-RhoA^G17A^ beads were added in each lysates and samples were rotated for 45 minutes at 4 °C. After washing the beads three times with Rho GEF buffer, the supernatants were carefully aspirated and additional 10 μl of Rho GEF buffer was added. Then, samples were denatured in 2x Laemmli buffer mixed with 2- mercaptoethanol by boiling at 75 °C for 15 minutes and SDS-PAGE analysis was performed as described.

### Immunostaining

We used collagen rat tail 1 (Gibco) 150 μg/ml for coating nano-patterned substrate (width: 800 nm, height: 800 nm and space: 800 nm) and 30 μg/ml for coating cover glass slides, both for 2 hours at room temperature. Cells were washed three times with PBS with MgCl2 1mM, CaCl2 1mM and incubated with 4 % paraformaldehyde (PFA) for 15 minutes at room temperature. Fixed cells were permeabilized with 0.1 % Triton X-100 in PBS for 5 minutes and incubated in blocking solution (10 % goat serum in PBS) for 1 hour at room temperature, followed by an incubation with primary antibody overnight at 4 °C. Next day, samples were washed with PBS three times and treated with secondary antibody for 1 hour at room temperature. After washing with PBS three times, samples were applied with ProLong Glass Antifade Mountant (Invitrogen, P36982). For imaging fixed samples, we used a laser scanning confocal microscope (Leica SP8, Yale West Campus Imaging Core) and an epi-fluorescent microscope (Zeiss Axiovert 200M).

### Cell transfection and virus infection

QIAprep Spin Miniprep kit (QIAGEN) was used for purifying plasmids. Transfections were carried out using Turbofect (Thermo Scientific) according to manufacturer’s recommendations. HEK 293T cells were cultured in T-75 flask and transfected at 70 – 90 % cell confluency with psPAX2 4 μg, pMD2.G 2 μg, target vector 4 μg or MLV gag/pol 4 μg, VSVG 2 μg, pCB150-2_GFP-EB1 4 μg for retrovirus production (day 0). Next day, we refreshed cell culture media and collected supernatants twice a day (morning and late afternoon) at day 2 and day 3. Then, the collected supernatants were mixed in PEG solution (final concentration 0.1 g/ml PEG 6000, NaCl 0.3M in deionized water, autoclaved) and stored at 4 °C overnight more than 12 hours. The supernatant/PEG solution mixture were centrifuged at 1500 × g for 30 minutes at 4 °C. After carefully removing supernatants from the tube, 1/10 of pellet volume DMEM with 25 mM HEPES was added and stored at −80 °C. On the day of infection, we injected with 5 μl/ml concentrated virus with polybrene 10 μg/ml to 70 – 90 % confluent cells. We cultured cells with virus for two days and began antibiotic selection with puromycin 1 μg/ml or blasticidin 5 μg/ml. For F-TRActin eGFP transfection, cells were transfected using FuGENE^®^HD Transfection reagents where 2 μg of the DNA plasmid was added to the transfection reagent and added to a cell dish. Cells were incubated for over 24 hours with the plasmid to complete the transfection process. Following 1 week of incubation with a selection media containing G418 (Mirus Bio LLC) at 1 mg/mL, population of cells were isolated and flow sorted with FACS flow cytometer. The isolated population were cultured and expanded DMEM (supplemented with 10 % FBS, 1 % Penstrep) and used for experimental purposes.

A construct of pCMV-GCaMP5G (Douglas Kim & Loren Looger from Addgene plasmid # 31788) was performed with a polymerase chain reaction and flanked with BamHI and XbaI site, then put into pLenti-mp2 backbone (Pantelis Tsoulfas from Addgene plasmid # 36097).

A construct of GEF-H1-eGFP was a kind gift from Robert Rottapel (University of Toronto, Ontario, Canada). pEGFP-C1 was obtained from clontech. GST – RhoA G17A, c-Flag pcDNA3 and pCB150-2_GFP-EB1 constructs were kind gifts from Rafael Garcia-Mata, Stephen Smale and Iain Cheeseman, respectively (Addgene plasmid #69357, #20011 and #46364). F-TRActin eGFP construct was a kind gift from Sergey Plotnikov (University of Toronto, Ontario, Canada). Scrambled shRNA plasmid was a kind gift from David Sabatini (Addgene plasmid #1864). shRNA plasmid against human GEF-H1 was obtained from Sigma (clone ID: NM_004723.x-235s1c1, sequence: CCGGCCCAACCTGCAATGTGACTATCTCGAGATAGTCACATTGCAGGTTGGGTTTTT).

### Real Time Polymerase Chain Reaction

RNA purification was carried out using RNeasy Plus Mini kit (QIAGEN) according to manufacturer’s recommendation. iScript Reverse Transcription Supermix (BioRad) was used for reverse transcription reaction with the suggested protocol (priming for 5 mins at 25 °C, reverse transcription for 20 mins at 46 °C and inactivation for 1 min at 95 °C) in a thermal cycler (C1000 Touch, Biorad). TaqMan Fast Advanced Master Mix (Applied Biosystems) was used for gene expression assays with the suggested protocol (incubation for 2 mins at 50 °C, polymerase activation for 20 secs at 95 °C, 40 cycles of PCR denaturing for 5 secs at 95 °C and annealing/extending for 30 secs at 55 °C) in real-time PCR system (CFX384 Touch, Biorad).

The following Taqman primers were used and obtained from Applied Biosystems: ARHGEF2 (Hs01064532_m1), GAPDH (Hs02786624_g1). All results were normalized to GAPDH from the same well. Each experiment was carried out in triplicate from three independent biological replicates.

### Live cell imaging

We autoclaved PDMS devices and sterilized glass slides (2460-1, Fisher Scientific) with 100 % EtOH for 30 minutes beforehand. In some experiments including collective cell migration, we used 35 mm dish with 20 mm glass bottom well (Cellvis). Both dried sides of PDMS and glass substrates underwent with plasma cleaning by hand-held corona treater (Electro-Technic, BD- 20) for 2 minutes, followed by an oven incubation at 75 °C for 30 minutes to facilitate tight bonding. We coated glass substrates with collagen rat tail 1 (Gibco) 30 μg/ml for 2 hours, washed with PBS three times before introducing cells. For spatially confining channel experiments, 8 × 10^6^ cells/ml were introduced from cell inlet and cultured > 15 hours before mounting on the scope.

Live cell imaging for FRET and Ca^2+^ experiments were performed on an epi-fluorescent microscope stage (Zeiss Axiovert 200M) with an environmental chamber set at 37 °C and 5 % CO_2_, coupled to a Cascade 512B II CCD (charge-coupled device) camera. The system was run by Slidebook software ver.6 (Intelligent Imaging Innovations) for automated control. All FRET experiments were captured every 5 minutes over at least 5-hour time period.

EB1 tracking and actin visualization experiments were performed on a spinning disk confocal microscope stage (Nikon Ti-E, Yale West Campus Imaging Core) with an environmental chamber set at 37 °C and 5 % CO_2_, coupled to Andor Zyila CMOS camera with 100x oil objective lens.

### Image analysis

Cells located under unconfined area at initial time point (time 0) were selected for Ca^2+^ activity analysis. Cell boundaries were drawn manually based on the GFP images and pixel intensities were measured using Fiji. The fluorescence intensity in cells in the spatially confining channels was normalized by the intensity at time 0.

The following image analysis was carried out using custom scripts written in MATLAB 2018b (MathWorks). For FRET analysis, fluorescence images were analyzed to background-correct images by subtracting the fluorescence intensity of background with no cells from the emission intensities of cells expressing fluorescent reporters. Background subtracted images were thresholded using Otsu’s method, registered, and then divided to produce ratiometric (FRET/CFP) images. To investigate the speed of cells, the position of cells at each time points was generated by obtaining centroids from the boundary of each cell based on the phase images. For visualization, after subtracting background images from fluorescent images, we applied thresholding using ‘adapthisteq’ function with 5x pixel intensity values. Then, images were registered to line up pixels, accounting for x-y translation of images then normalized to produce FRET/CFP images, then followed by pixel-by-pixel normalization to points indicated by experiments.

Peaks of pulsatile time-dependent functions were determined using the algorithm developed by Tom O’Haver (the University of Maryland at College Park). The code searches for downward zero crossings in the first derivative of the power spectrum. The same parameter values (Slope Threshold: < 0.0001, Smooth Width: 4, 5 and Fit Width: 12) were used to determine the number of peaks for all experiments. (https://terpconnect.umd.edu/~toh/spectrum/PeakFindingandMeasurement.htm#ipeak)

To track EB1 proteins and evaluate microtubule dynamics parameters, we used a particle tracking MATLAB function written by Daniel Blair (Georgetown University) and Eric Dufresne (ETH Zürich). Briefly, the function records the localization of EB1 proteins based on the GFP intensity after smoothing images by spatial bandpass filter and subtracting the background off. Then, the location data are tracked between frames to calculate the speed and the angle of EB1 spot trajectories using custom written scripts. Threshold value of EB1 protein was determined using the Otsu’s method. The diameter of the EB1 spot was directly measured in pixels. We used the same image analysis parameters for all experiments. After calculating speeds of EB1 proteins over 2-minute time period, two ends of standard deviation (sigma) were excluded for final speed calculation in order to remove outliers. The original particle tracking function is available at the following link. (https://site.physics.georgetown.edu/matlab/tutorial.html)

### Polyacrylamide gel formation preparation

Traction force Microscopy experiments were carried out on polyacrylamide (PA) gels, polymerized onto 25 mm diameter (#1.5, Dow Corning) coverslips. Briefly, the coverslips are treated with a combination of aminopropylsilane (Sigma Aldrich) and glutaraldehyde (Electron Microscopy Sciences) to make the surface reactive to the acrylamide. The ratios of polyacrylamide to bis-acrylamide for the gels is used as 7.5 %: 0.153 % to yield a gel with an elastic modulus of E= 4.3 kPa. A concentration of 0.05 % w/v ammonium persulfate (Fisher BioReagents) and 20 nM beads (Molecular Probes) of 0.1 μm size are embedded in the gel mixture prior to polymerization. A 15 μl of the gel is added to the coverslip and covered with another coverslip, which has been made hydrophobic through treatment with Rain-X^®^. The gels are polymerized on the coverslips for 30 minutes at room temperature. The gels are then reacted with the standard 1 mg/mL Sulfo-SANPAH (Thermo Fisher Scientific). The surface of the gels is then coated with fibronectin (F002, Sigma-Aldrich) with 0.5 mg/ml concentration. The reaction proceeds for 12 hours overnight incubation in the dark, and the coverslips are then rinsed and stored in 1X PBS.

### Traction force measurement

Traction force microscopy is used to measure the forces exerted by cells on the substrate using confocal imaging. Cells were plated for 12 hours on 4.3 kPa PA gels before imaging. ‘Force-loaded’ images (with cells) of the beads embedded in the polyacrylamide gels were obtained using a 40X oil-immersion objective (Leica Microsystems). The ‘null-force’ image was obtained at the end of each experiment by adding trypsin to the cells for 1 hour. Confocal bead images were aligned to correct for drift (StackReg in ImageJ) and compared to the reference image using PIV software (http://www.oceanwave.jp/softwares/mpiv/) to obtain the displacements field (**x**). Forces were calculated using custom-written code by Ulrich Schwarz (Sabass et al., 2008). The traction forces were used to calculate the energies of deformation (strain energies) in the cell adhesion substratum using the spatial distribution of forces **F** and displacements **x**. The spatial distribution of strain energies is calculated, for each coarse-grained grid size, using 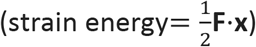. The total strain energy for a given cell area is calculated by summing up the local values within each grid.

## Acknowledgments

The work was supported by NIH grants CA155758-05 (to S.H.L., A.H., H.C. and A.L.) and U54 CA209992 (to S.H.L., A.H., S.Y.M., V.A., H.C., M.M. and A.L.). We thank Andrew Ewald and Alexander Popel for helpful comments.

## Author Contributions

S.H.L. and A.L. conceived the project. S.H.L. designed and used the 3D collective migration device, performed all experiments unless otherwise indicated and analyzed the data. A.H. analyzed the FRET data. M.S.Y. performed the traction force experiment and analysis of the data. V.A. and M.S.Y. fabricated the devices for the analysis of cell migration in spatially confining channels. H.C. constructed the plasmid for detecting Ca^2+^ activity (GCaMP5). S.H.L. and A.L. performed the computational modeling analysis. S.H.L. and A.L. wrote and edited the manuscript. M.M. and A.L. obtained the funding. A.L. supervised the project.

## References

Abrams, G.A., Bentley, E., Nealey, P.F., and Murphy, C.J. (2002). Electron microscopy of the canine corneal basement membranes. Cells Tissues Organs 170, 251–257.

Bartolini, F., Andres-Delgado, L., Qu, X.Y., Nik, S., Ramalingam, N., Kremer, L., Alonso, M.A., and Gundersen, G.G. (2016). An mDia1-INF2 formin activation cascade facilitated by IQGAP1 regulates stable microtubules in migrating cells. Mol Biol Cell 27, 1797–1808.

Belotti, D., Rieppi, M., Nicoletti, M.I., Casazza, A.M., Fojo, T., Taraboletti, G., and Giavazzi, R. (1996). Paclitaxel (Taxol(R)) inhibits motility of paclitaxel-resistant human ovarian carcinoma cells. Clin Cancer Res 2, 1725–1730.

Bijman, M.N.A., Amerongen, G.P., Laurens, N., van Hinsbergh, V.W.M., and Boven, E. (2006). Microtubule-targeting agents inhibit angiogenesis at subtoxic concentrations, a process associated with inhibition of Rac1 and Cdc42 activity and changes in the endothelial cytoskeleton. Mol Cancer Ther 5, 2348–2357.

Bouchet, B.P., and Akhmanova, A. (2017). Microtubules in 3D cell motility. J Cell Sci 130, 39–50.

Bowers, E.M., Yan, G., Mukherjee, C., Orry, A., Wang, L., Holbert, M.A., Crump, N.T., Hazzalin, C.A., Liszczak, G., Yuan, H., et al. (2010). Virtual ligand screening of the p300/CBP histone acetyltransferase: identification of a selective small molecule inhibitor. Chem Biol 17, 471–482.

Chauhan, B.K., Lou, M., Zheng, Y., and Lang, R.A. (2011). Balanced Rac1 and RhoA activities regulate cell shape and drive invagination morphogenesis in epithelia. Proc Natl Acad Sci U S A 108, 18289–18294.

Chen, X., Wanggou, S., Bodalia, A., Zhu, M., Dong, W., Fan, J.J., Yin, W.C., Min, H.K., Hu, M., Draghici, D., et al. (2018). A Feedforward Mechanism Mediated by Mechanosensitive Ion Channel PIEZO1 and Tissue Mechanics Promotes Glioma Aggression. Neuron 100, 799–815 e797.

Cheong, R., and Levchenko, A. (2010). Oscillatory signaling processes: the how, the why and the where. Curr Opin Genet Dev 20, 665–669.

Devreotes, P., and Horwitz, A.R. (2015). Signaling Networks that Regulate Cell Migration. Cold Spring Harb Perspect Biol 7.

Dogterom, M., and Koenderink, G.H. (2019). Actin-microtubule crosstalk in cell biology. Nat Rev Mol Cell Biol 20, 38–54.

Dolmetsch, R.E., Xu, K.L., and Lewis, R.S. (1998). Calcium oscillations increase the efficiency and specificity of gene expression. Nature 392, 933–936.

Doyle, A.D., Petrie, R.J., Kutys, M.L., and Yamada, K.M. (2013). Dimensions in cell migration. Curr Opin Cell Biol 25, 642–649.

Doyle, A.D., Wang, F.W., Matsumoto, K., and Yamada, K.M. (2009). One-dimensional topography underlies three-dimensional fibrillar cell migration. J Cell Biol 184, 481–490.

Endo, M., Nishita, M., Fujii, M., and Minami, Y. (2015). Insight into the role of Wnt5a-induced signaling in normal and cancer cells. Int Rev Cell Mol Biol 314, 117–148.

Fish, J.E., Gutierrez, M.C., Dang, L.T., Khyzha, N., Chen, Z.Q., Veitch, S., Cheng, H.S., Khor, M., Antounians, L., Njock, M.S., et al. (2017). Dynamic regulation of VEGF-inducible genes by an ERK/ERG/p300 transcriptional network. Development 144, 2428–2444.

Friedl, P., and Alexander, S. (2011). Cancer invasion and the microenvironment: plasticity and reciprocity. Cell 147, 992–1009.

Friedl, P., and Wolf, K. (2010). Plasticity of cell migration: a multiscale tuning model. J Cell Biol 188, 11–19.

Garcia-Mata, R., Wennerberg, K., Arthur, W.T., Noren, N.K., Ellerbroek, S.M., and Burridge, K. (2006). Analysis of activated GAPs and GEFs in cell lysates. Methods in Enzymology, Vol 406, Regulators and Effectors of Small Gtpases: Rho Family 406, 425–437.

Geudens, I., and Gerhardt, H. (2011). Coordinating cell behaviour during blood vessel formation. Development 138, 4569–4583.

Guilluy, C., Dubash, A.D., and Garcia-Mata, R. (2011). Analysis of RhoA and Rho GEF activity in whole cells and the cell nucleus. Nat Protoc 6, 2050–2060.

Gundersen, G.G., and Bulinski, J.C. (1988). Selective stabilization of microtubules oriented toward the direction of cell migration. Proc Natl Acad Sci U S A 85, 5946–5950.

Hind, L.E., Vincent, W.J., and Huttenlocher, A. (2016). Leading from the Back: The Role of the Uropod in Neutrophil Polarization and Migration. Dev Cell 38, 161–169.

Hirokawa, N., Noda, Y., and Okada, Y. (1998). Kinesin and dynein superfamily proteins in organelle transport and cell division. Curr Opin Cell Biol 10, 60–73.

Hung, W.C., Chen, S.H., Paul, C.D., Stroka, K.M., Lo, Y.C., Yang, J.T., and Konstantopoulos, K. (2013). Distinct signaling mechanisms regulate migration in unconfined versus confined spaces. J Cell Biol 202, 807–824.

Hung, W.C., Yang, J.R., Yankaskas, C.L., Wong, B.S., Wu, P.H., Pardo-Pastor, C., Serra, S.A., Chiang, M.J., Gu, Z.Z., Wirtz, D., et al. (2016). Confinement Sensing and Signal Optimization via Piezo1/PKA and Myosin II Pathways. Cell Rep 15, 1430–1441.

Kim, D.H., Han, K., Gupta, K., Kwon, K.W., Suh, K.Y., and Levchenko, A. (2009). Mechanosensitivity of fibroblast cell shape and movement to anisotropic substratum topography gradients. Biomaterials 30, 5433–5444.

Kim, D.H., Provenzano, P.P., Smith, C.L., and Levchenko, A. (2012). Matrix nanotopography as a regulator of cell function. J Cell Biol 197, 351–360.

Knofler, M., and Pollheimer, J. (2013). Human placental trophoblast invasion and differentiation: a particular focus on Wnt signaling. Front Genet 4, 190.

Kolovos, P., Georgomanolis, T., Koeferle, A., Larkin, J.D., Brant, L., Nikolic, M., Gusmao, E.G., Zirkel, A., Knoch, T.A., van Ijcken, W.F., et al. (2016). Binding of nuclear factor kB to noncanonical consensus sites reveals its multimodal role during the early inflammatory response. Genome Res 26, 1478–1489.

Krendel, M., Zenke, F.T., and Bokoch, G.M. (2002). Nucleotide exchange factor GEF-H1 mediates cross-talk between microtubules and the actin cytoskeleton. Nat Cell Biol 4, 294–301.

Lasko, L.M., Jakob, C.G., Edalji, R.P., Qiu, W., Montgomery, D., Digiammarino, E.L., Hansen, T.M., Risi, R.M., Frey, R., Manaves, V., et al. (2017). Discovery of a selective catalytic p300/CBP inhibitor that targets lineage-specific tumours. Nature 550, 128–132.

Lawson, C.D., and Ridley, A.J. (2018). Rho GTPase signaling complexes in cell migration and invasion. J Cell Biol 217, 447–457.

Liu, Y.J., Le Berre, M., Lautenschlaeger, F., Maiuri, P., Callan-Jones, A., Heuze, M., Takaki, T., Voituriez, R., and Piel, M. (2015). Confinement and Low Adhesion Induce Fast Amoeboid Migration of Slow Mesenchymal Cells. Cell 160, 659–672.

Mackeh, R., Lorin, S., Ratier, A., Mejdoubi-Charef, N., Baillet, A., Bruneel, A., Hamai, A., Codogno, P., Pous, C., and Perdiz, D. (2014). Reactive oxygen species, AMP-activated protein kinase, and the transcription cofactor p300 regulate alpha-tubulin acetyltransferase-1 (aTAT-1/MEC-17)-dependent microtubule hyperacetylation during cell stress. J Biol Chem 289, 11816–11828.

Mao, Z.X., Bonni, A., Xia, F., Nadal-Vicens, M., and Greenberg, M.E. (1999). Neuronal activity-dependent cell survival mediated by transcription factor MEF2. Science 286, 785–790.

Maruta, H., Greer, K., and Rosenbaum, J.L. (1986). The Acetylation of Alpha-Tubulin and Its Relationship to the Assembly and Disassembly of Microtubules. J Cell Biol 103, 571–579.

Miyamoto, T., Mochizuki, T., Nakagomi, H., Kira, S., Watanabe, M., Takayama, Y., Suzuki, Y., Koizumi, S., Takeda, M., and Tominaga, M. (2014). Functional Role for Piezo1 in Stretch-evoked Ca^2+^ Influx and ATP Release in Urothelial Cell Cultures. J Biol Chem 289, 16565–16575.

Munevar, S., Wang, Y.L., and Dembo, M. (2001). Traction force microscopy of migrating normal and H-ras transformed 3T3 fibroblasts. Biophys J 80, 1744–1757.

Noren, D.P., Chou, W.H., Lee, S.H., Qutub, A.A., Warmflash, A., Wagner, D.S., Popel, A.S., and Levchenko, A. (2016). Endothelial cells decode VEGF-mediated Ca^2+^ signaling patterns to produce distinct functional responses. Science Signaling 9, ra20.

O’Connell, M.P., Marchbank, K., Webster, M.R., Valiga, A.A., Kaur, A., Vultur, A., Li, L., Herlyn, M., Villanueva, J., Liu, Q., et al. (2013). Hypoxia induces phenotypic plasticity and therapy resistance in melanoma via the tyrosine kinase receptors ROR1 and ROR2. Cancer Discov 3, 1378–1393.

Palazzo, A.F., Cook, T.A., Alberts, A.S., and Gundersen, G.G. (2001). mDia mediates Rho-regulated formation and orientation of stable microtubules. Nat Cell Biol 3, 723–729.

Paluch, E.K., Aspalter, I.M., and Sixt, M. (2016). Focal Adhesion-Independent Cell Migration. Annu Rev Cell Dev Biol 32, 469–490.

Pardo-Pastor, C., Rubio-Moscardo, F., Vogel-Gonzalez, M., Serra, S.A., Afthinos, A., Mrkonjic, S., Destaing, O., Abenza, J.F., Fernandez-Fernandez, J.M., Trepat, X., et al. (2018). Piezo2 channel regulates RhoA and actin cytoskeleton to promote cell mechanobiological responses. Proc Natl Acad Sci U S A 115, 1925–1930.

Park, J., Holmes, W.R., Lee, S.H., Kim, H.N., Kim, D.H., Kwak, M.K., Wang, C.J., Edelstein-Keshet, L., and Levchenko, A. (2017). Mechanochemical feedback underlies coexistence of qualitatively distinct cell polarity patterns within diverse cell populations. Proc Natl Acad Sci U S A 114, E5750–E5759.

Park, J., Kim, D.H., Shah, S.R., Kim, H.N., Kshitiz, Kim, P., Quinones-Hinojosa, A., and Levchenko, A. (2019). Switch-like enhancement of epithelial-mesenchymal transition by YAP through feedback regulation of WT1 and Rho-family GTPases. Nat Commun 10, 2797.

Petrie, R.J., Doyle, A.D., and Yamada, K.M. (2009). Random versus directionally persistent cell migration. Nat Rev Mol Cell Bio 10, 538–549.

Prahl, L.S., Bangasser, P.F., Stopfer, L.E., Hemmat, M., White, F.M., Rosenfeld, S.S., and Odde, D.J. (2018). Microtubule-Based Control of Motor-Clutch System Mechanics in Glioma Cell Migration. Cell Rep 25, 2591–2604.

Ridley, A.J., Schwartz, M.A., Burridge, K., Firtel, R.A., Ginsberg, M.H., Borisy, G., Parsons, J.T., and Horwitz, A.R. (2003). Cell migration: integrating signals from front to back. Science 302, 1704–1709.

Sabass, B., Gardel, M.L., Waterman, C.M., and Schwarz, U.S. (2008). High resolution traction force microscopy based on experimental and computational advances. Biophys J 94, 207–220.

Sacilotto, N., Chouliaras, K.M., Nikitenko, L.L., Lu, Y.W., Fritzsche, M., Wallace, M.D., Nornes, S., Garcia-Moreno, F., Payne, S., Bridges, E., et al. (2016). MEF2 transcription factors are key regulators of sprouting angiogenesis. Gene Dev 30, 2297–2309.

Sheng, M., Thompson, M.A., and Greenberg, M.E. (1991). Creb – a Ca^2+^-Regulated Transcription Factor Phosphorylated by Calmodulin-Dependent Kinases. Science 252, 1427–1430.

Siegrist, S.E., and Doe, C.Q. (2007). Microtubule-induced cortical cell polarity. Genes Dev 21, 483–496.

Tabdanov, E.D., Puram, V., Zhovmer, A., and Provenzano, P.P. (2018). Microtubule-Actomyosin Mechanical Cooperation during Contact Guidance Sensing. Cell Rep 25, 328–338 e325.

Watanabe, N., and Higashida, C. (2004). Formins: processive cappers of growing actin filaments. Exp Cell Res 301, 16–22.

Weisenberg, R.C. (1972). Microtubule Formation in-Vitro in Solutions Containing Low Calcium Concentrations. Science 177, 1104–1105.

Weisenberg, R.C., and Deery, W.J. (1981). The Mechanism of Calcium-Induced Microtubule Disassembly. Biochem Biophys Res Commun 102, 924–931.

Yang, H.L., Ganguly, A., and Cabral, F. (2010). Inhibition of Cell Migration and Cell Division Correlates with Distinct Effects of Microtubule Inhibiting Drugs. J Biol Chem 285, 32242–32250.

Yokota, Y., Nakajima, H., Wakayama, Y., Muto, A., Kawakami, K., Fukuhara, S., and Mochizuki, N. (2015). Endothelial Ca^2+^ oscillations reflect VEGFR signaling-regulated angiogenic capacity in vivo. Elife 4, e08817.

Youn, H.D., Sun, L., Prywes, R., and Liu, J.O. (1999). Apoptosis of T cells mediated by Ca^2+^-induced release of the transcription factor MEF2. Science 286, 790–793.

Zhang, J., Guo, W.H., and Wang, Y.L. (2014). Microtubules stabilize cell polarity by localizing rear signals. Proc Natl Acad Sci U S A 111, 16383–16388.

