## Supplementary Figures for "A molecular clock controls periodically driven cell migration in confined spaces"

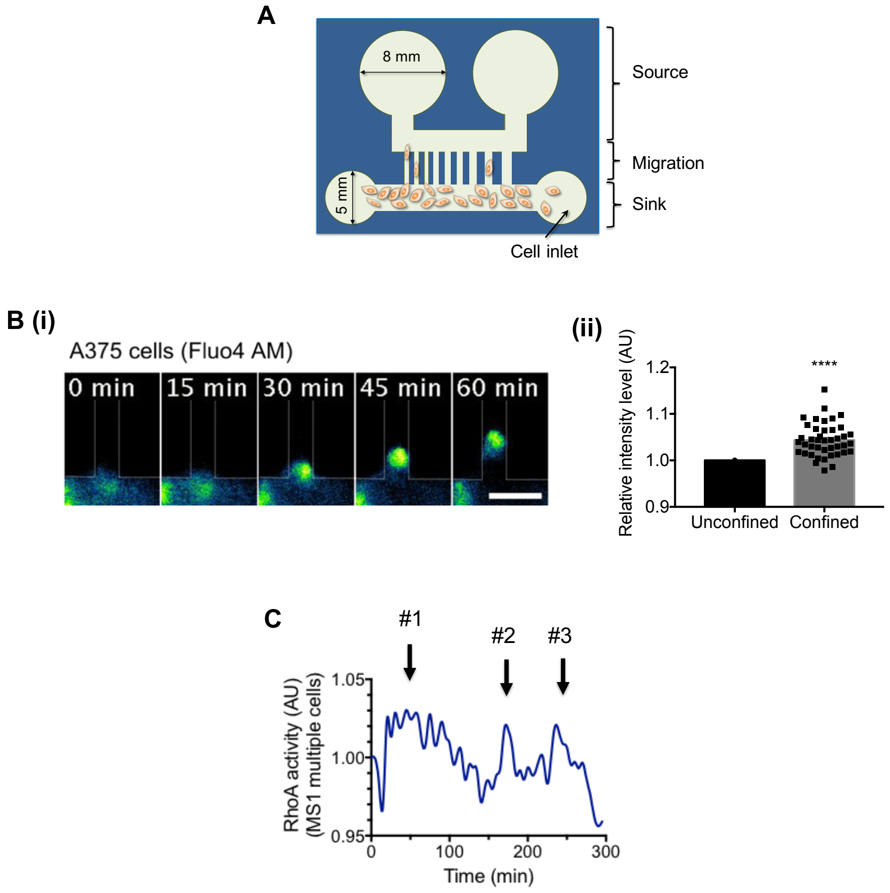


**Supplementary Figure 1. Further characterization of cell migration under spatial confinement**
**(A)** Schematic of the microfluidic device used to generate physical confinements (width: 3, 6, 10, 20 and 50 μm, height: 10 μm and length: 200 μm); **(B)** (i) Representative time-lapse images of Fluo4 AM dyed A375 cells migrating in channel (version1). Scale bar: 10 μm. (ii) Comparison of relative fluorescence intensity levels of cells before (unconfined) and after (confined) entering the channel. Data were analyzed by unpaired two-tailed t test with error bar representing s.e.m. (n = 41 cells, **** $P < 0.0001$); **(C)** RhoA activity of multiple MS1 cells migrating under spatial confinement.

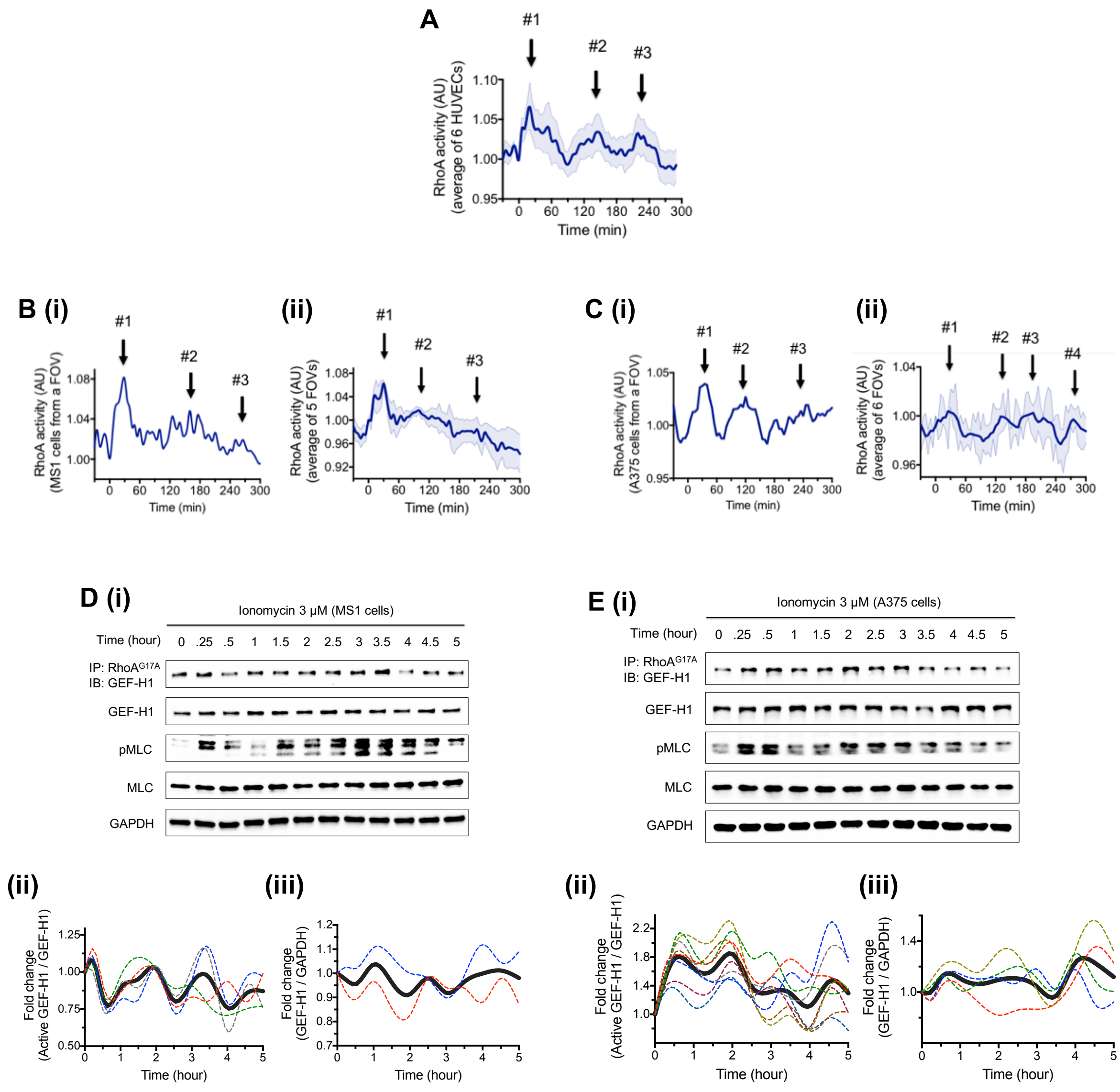


**Supplementary Figure 2. Further characterization of cell responses to a persistent increase in the intracellular $Ca^{2+}$ concentration in the absence of spatial confinement. (A)** Average RhoA activity from 6 different HUVECs with error bar representing s.e.m. **(B)** (i) Average RhoA activity of MS1 cells from a single field of view (FOV) (n=17 cells). (ii) Average RhoA activity of MS1 cells from 5 different FOVs with error bar representing s.e.m. Each FOV, obtained by 40x objective lens, contained an average of 18 cells. **(C)** (i) Average RhoA activity of A375 cells from a single FOV (n=42 cells). (ii) Average RhoA activity of A375 cells from 6 different FOVs with error bar representing s.e.m. Each FOV, obtained by 40x objective lens, contained an average of 39 cells. Graph was processed with 2nd order smoothing with 4 neighbors using the function in Prism7 for (C). **(D,E)** Immunoblot analysis of the activation level and total abundance of GEF-H1, and of pMLC and MLC in (D) (i) MS1 and (E) (i) A375 cells in response to treatment with ionomycin (3 μM). GAPDH was used as a loading control. (D,E) (ii) Fold-change quantification of the active GEF-H1 level normalized over the total GEF-H1 abundance and (iii) total GEF-H1 level normalized over GAPDH. The dotted colored curves indicate results of independent experiments and the solid gray curve represents the average of dotted lines. The curves were generated by the method of cubic spline interpolation using the function in Prism7.

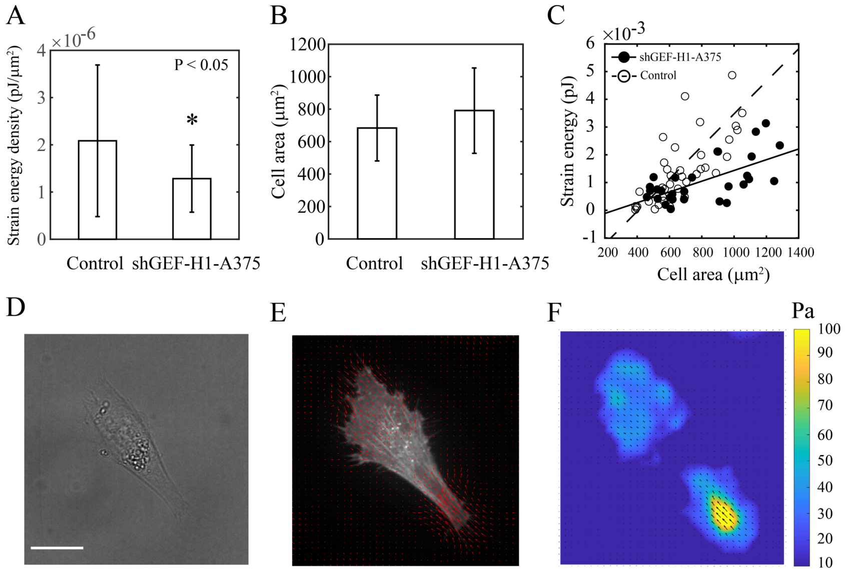


**Supplementary Figure 3. Traction force measurement in shGEF-H1 A375 cells**

**(A)** Strain energy density and **(B)** cell spread area on 4.3kPa polyacrylamide adhesion substratum **(C)** Strain energy increases linearly with the cell spreading area for control and shGEF-H1-A375 cells. **(D)** DIC image. **(E)** Traction force vectors superimposed on actin image for the cell shown in (D), **(F)** Stress map. Scale bar: 50 μm. (n=30 (control), n=25 (shGEF-H1 A375 cells)). See Methods for details.

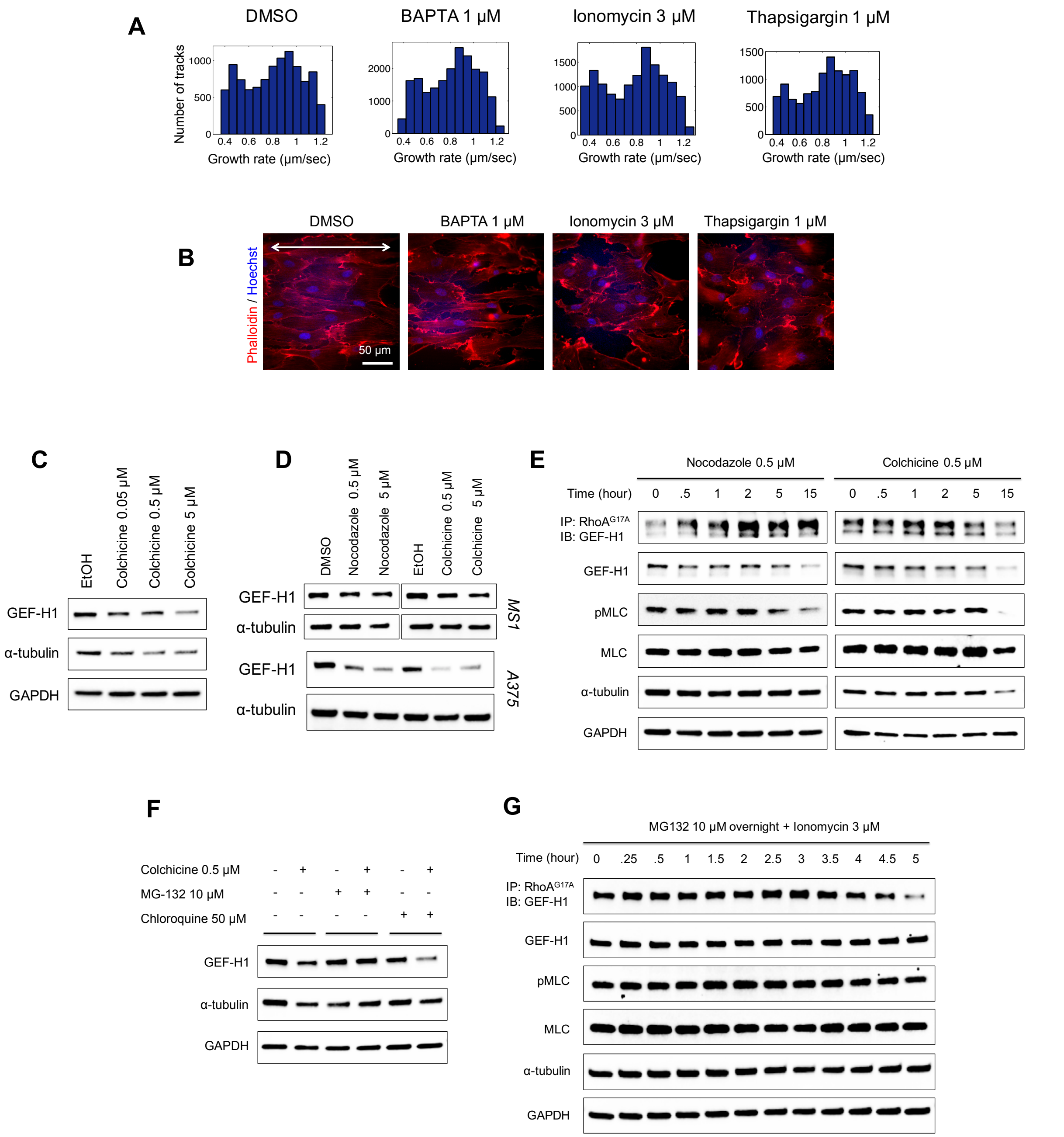


**Supplementary Figure 4. Additional analysis of the effect of $Ca^{2+}$ on MT stability and of MT stability on GEF-H1 degradation.** **(A)** The histograms of MT growth rate distributions corresponding to the experiment in Figure 3A (the results are summarized in Table S1). **(B)** Representative images of HUVECs stained with phalloidin (red) and Hoechst (blue) following treatment of DMSO, BAPTA (1 µM), ionomycin (3 µM) or thapsigargin (1 µM) for 15 hours. White arrow indicates the direction of the nano-scale ridges and grooves in the cell culture substratum. Scale bar: 50 µm. **(C)** Immunoblot of GEF-H1 and α-tubulin in HUVECs following treatment with the solvent (EtOH) and colchicine EtOH solution (0.05, 0.5, 5 µM) for 15 hours. GAPDH was used as a loading control. Analysis is based on 3 independent experiments. **(D)** Immunoblot of GEF-H1 abundance in MS1 and A375 cells following treatment of DMSO, nocodazole (0.5, 5 µM), EtOH and colchicine (0.5, 5 µM) for 15 hours. α-tubulin was used as a loading control. Analysis is based on 2 independent experiments. **(E)** Immunoblot of activation level and total abundance of GEF-H1, and of pMLC, MLC and α-tubulin in HUVECs following treatment with nocodazole (0.5 μM) or colchicine (0.5 μM). GAPDH was used as a loading control. Analysis is based on 2 independent experiments. **(F)** Immunoblot of GEF-H1 abundance and α-tubulin in HUVECs following treatment with MG-132 (10 μM), chloroquine (50 μM) and/or colchicine (0.5 μM) for 15 hours. Analysis is based on 3 independent experiments. **(G)** Immunoblot of activation level and total abundance of GEF-H1, and of pMLC, MLC and α-tubulin in HUVECs in response to treatment with ionomycin (3 μM) following pre-treatment with MG-132 (10 μM) for 15 hours. GAPDH was used as a loading control. Analysis is based on 2 independent experiments.

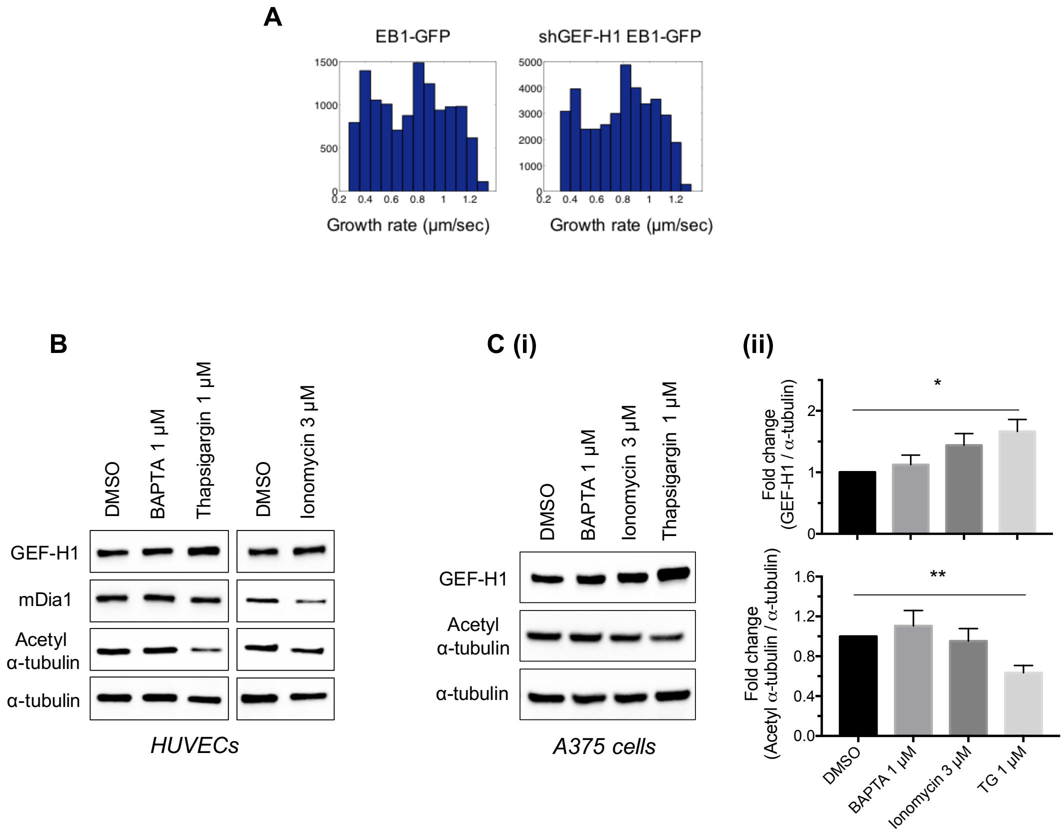


**Supplementary Figure 5. Further analysis of interplay between GEF-H1, mDia1 and microtubules.** **(A)** MT growth rate histograms of EB1-GFP in un-infected and shGEF-H1 expressing HUVECs. **(B)** Immunoblot analysis of the abundances of GEF-H1, mDia1 and acetylated α-tubulin in HUVECs following treatment of DMSO, BAPTA (1 μM), thapsigardin (1 μM) and ionomycin (3 μM) for 15 hours. α-tubulin was used as a loading control. Analysis is based on 2 independent experiments. **(C)** (i) Immunoblot analysis of the abundances of GEF-H1 and acetylated α-tubulin in A375 cells analyzed similarly to (B). α-tubulin was used as a loading control. (ii) Fold-change quantification of GEF-H1 abundance relative to α-tubulin (* P = 0.0274) and acetylated α-tubulin abundance relative α-tubulin (** P = 0.0079). Data were analyzed by unpaired two-tailed t test with error bar representing s.e.m. Analysis is based on 3 independent experiments.

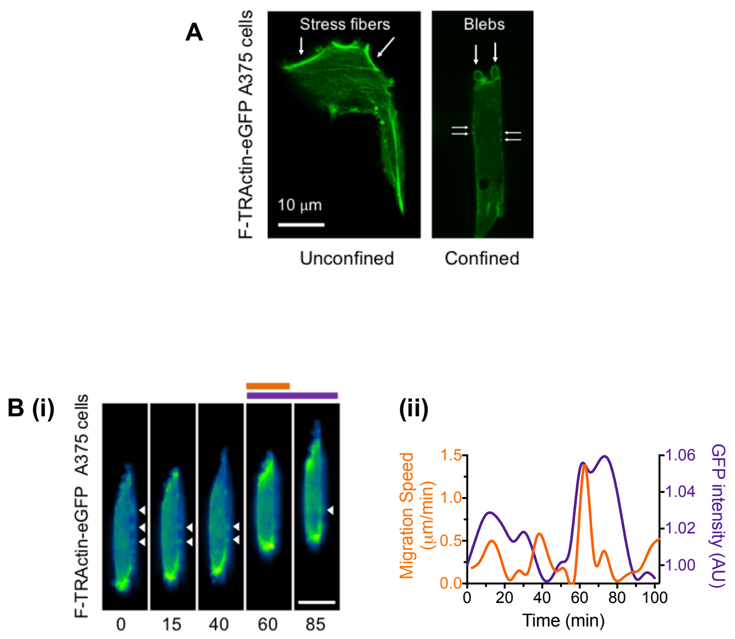


**Supplementary Figure 6. Further analysis of actin polymerization in migrating A375 cells**
**(A)** Representative images of F-TRActin-eGFP transfected A375 cells in unconfined (left) or spatially confining (right) spaces (version2). Scale bar: 10 µm. **(B)** (i) Time lapse images of F-TRActin-eGFP A375 cells migrating under spatial confinement for the condition as in Figure 6C. Horizontal bars indicate the timing of the migration speed (orange) and GFP intensity (purple) peaks corresponding to a portion of the graph below (bottom right). White arrows indicate blebs at the lateral cell periphery. Scale bar: 10 µm. (ii) Quantification of cell migration speed and the intensity of actin filamentation over time for the experiment in (i).

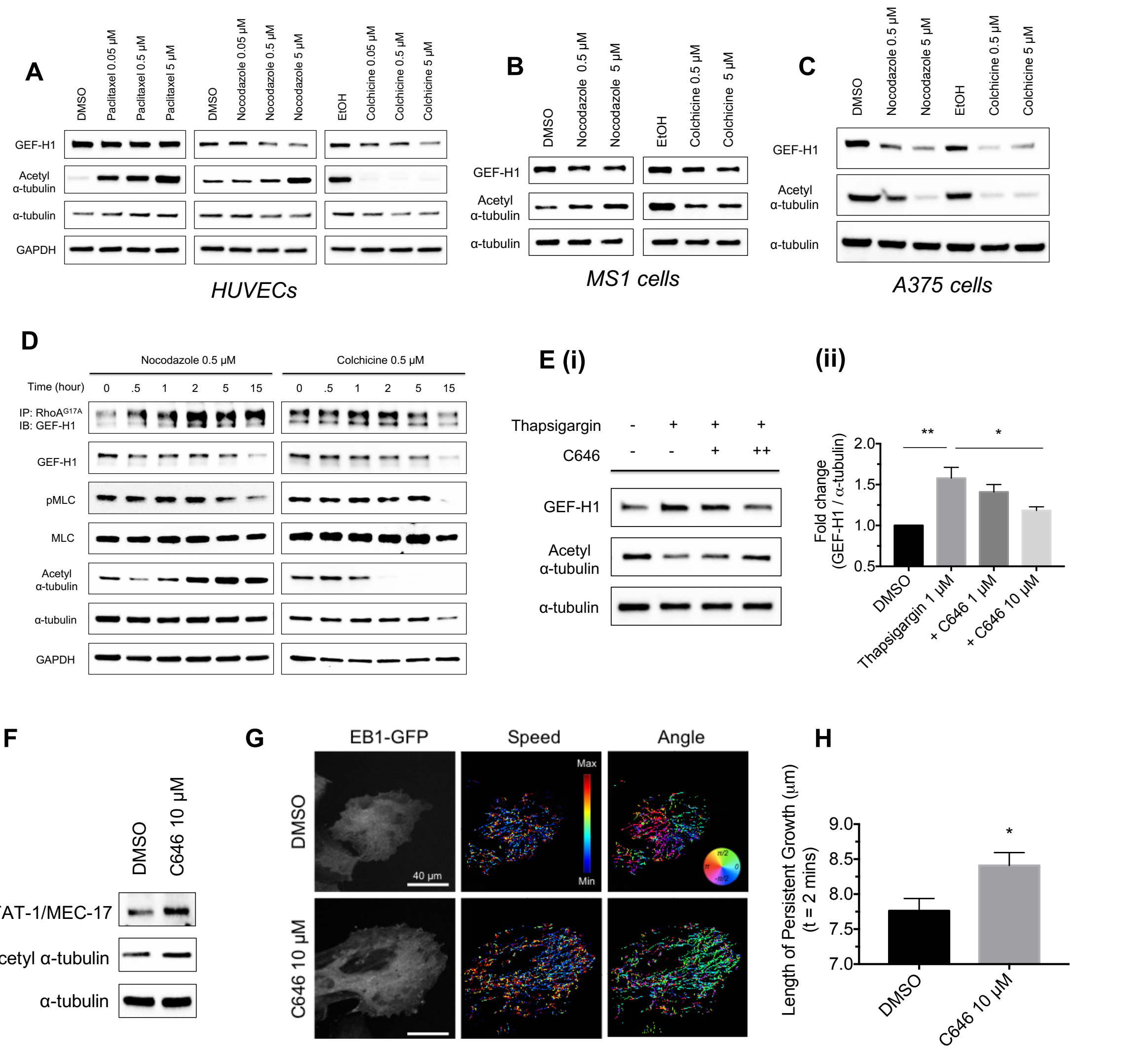


**Supplementary Figure 7. Tubulin acetylation is correlated with microtubule stability in endothelial and melanoma cells. (A-D)** Immunoblot analysis of acetylated α-tubulin abundance for experiments shown in in Figure 3D and Supplementary Figure 4C (A), Supplementary Figure 4D (B-C) and Supplementary Figure 4E (D). **(E)** (i) Immunoblot analysis of the abundances of GEF-H1 and acetylated α-tubulin in HUVECs following treatment with thapsigargin (1 μM) and/or C646 (1, 10 μM) for 15 hours. α-tubulin was used as a loading control. (ii) Fold-change quantification of GEF-H1 abundance relative to α-tubulin for the conditions in (i). Data were analyzed by one-way ANOVA with error bar representing s.e.m. (* P = 0.0256, ** P = 0.0018). Analysis is based on 4 independent experiments. **(F)** Immunoblot analysis of abundances of αTAT-1/MEC-17 and acetylated α-tubulin in HUVECs treated with DMSO or C646 (10 μM) for 15 hours. α-tubulin was used as a loading control. **(G)** Representative fluorescent images of HUVECs expressing EB1-GFP, treated with DMSO or C646 (10 μM) for 15 hours and the results of an automated tracking analysis representing speed and angle of EB1 trajectories from images taken every 2 seconds for 2 minutes. The analysis is performed as in Fig. 3. Scale bar: 40 μm. **(H)** Comparison of the length of persistent growth of microtubules over 2 minutes for cells treated with DMSO (n = 11 cells) or C646 (n = 13 cells) cells. Data were analyzed by unpaired two-tailed t test with error bar representing s.e.m. (* P = 0.0197)

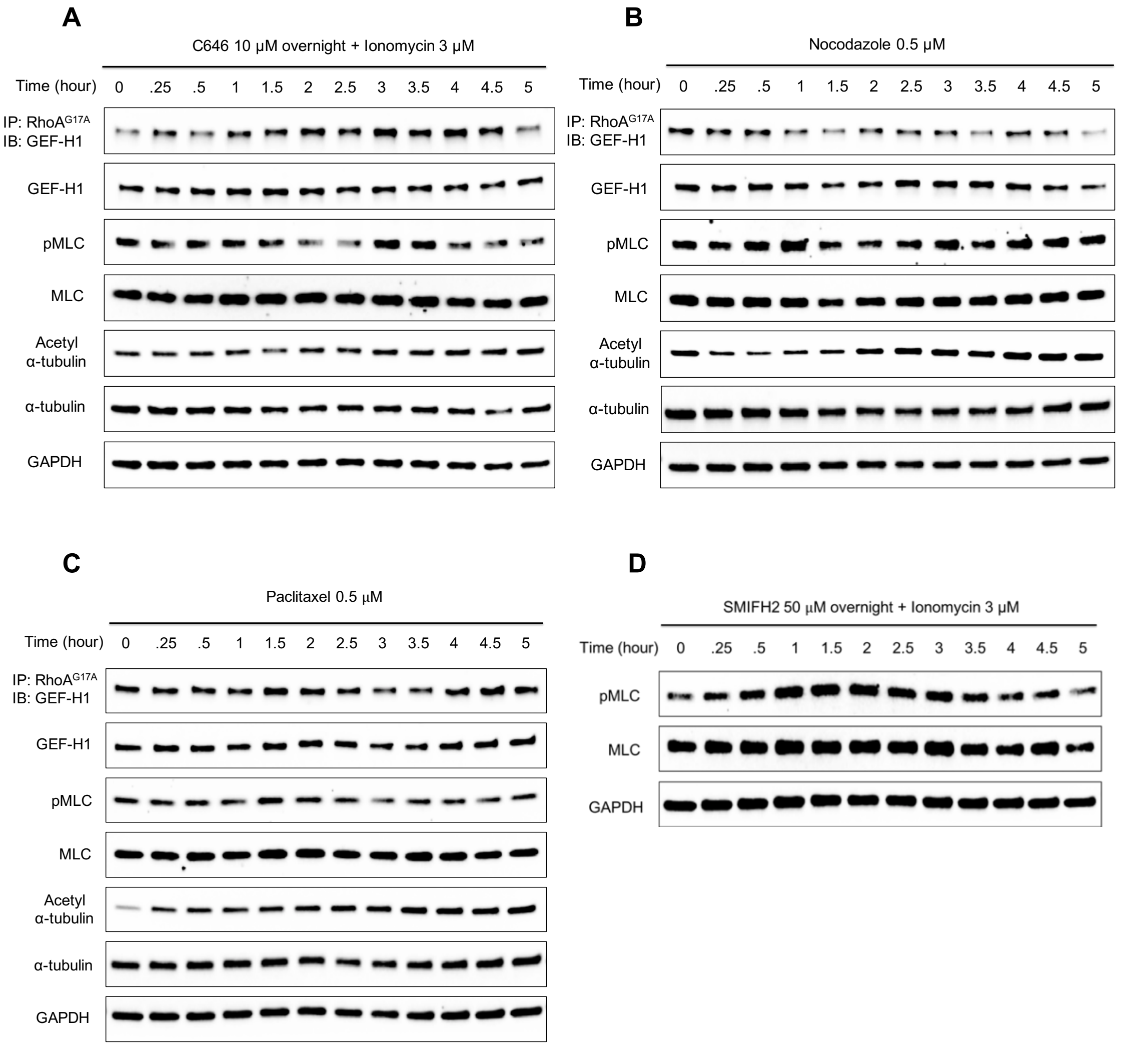


**Supplementary Figure 8.**
Immunoblot analysis of the activation level and total abundance of GEF-H1, and of pMLC, MLC, acetylated $\alpha$-tubulin and $\alpha$-tubulin in HUVECs in response to treatment with **(A)** ionomycin (3 $\mu$M) after pre-treatment of C646 (10 $\mu$M) for 15 hours, and treatment with **(B)** nocodazole (0.5 $\mu$M) and **(C)** paclitaxel (0.5 $\mu$M). **(D)** Immunoblot analysis of pMLC and MLC in HUVECs in response to treatment with ionomycin (3 $\mu$M) after pre-treatment of SMIFH2 (50 $\mu$M) for 15 hours. GAPDH was used as a loading control.

**Table S1.**

| | Control | BAPTA | Ionomycin | Thapsigargin |
|---|---|---|---|---|
| **Number of cells** | 11 | 14 | 14 | 14 |
| **Number of MTs tracked per cell** | 1151.7 | 1462.7 | 984.3 | 808.9 |
| **Growth rate (µm/sec)** | 0.787 ± 0.012 | 0.806 ± 0.01 | 0.764 ± 0.02 | 0.805 ± 0.013 |
| **Length of persistent growth (µm)** | 7.68 ± 0.334 | * 8.88 ± 0.411 | * 6.138 ± 0.42 | * 6.787 ± 0.264 |

**Table S2.**

| | EB1-GFP | shGEF-H1 EB1-GFP |
|---|---|---|
| **Number of cells** | 13 | 23 |
| **Number of MTs tracked per cell** | 937.6 | 1663.2 |
| **Growth rate (µm/sec)** | 0.756 ± 0.013 | 0.763 ± 0.015 |
| **Length of persistent growth (µm)** | 8.071 ± 0.151 | ** 8.945 ± 0.191 |
