## Supplementary Methods S1 for "A molecular clock controls periodically driven cell migration in confined spaces"

### Methods S1. Collective endothelial cell migration in 3D gel matrix

*In vivo*, physical cell confinement arises during migration in dense extracellular matrices (ECM). Although it is convenient to study cell migration in spatially confining spaces using microfabricated channels of defined dimensions, it is of interest to test whether the insights into the molecular mechanisms developed in such devices would translate into the more realistic 3D migration with the engineered or natural ECM. To develop a model allowing us to explore the migration of endothelial cells into 3D ECM micro-environment, mimicking the initial stages of angiogenesis, we sought to mimic not only invasive cell spread, but the initial separation of the 'Tip' cells from the monolayer of the parental blood vessel leading to growth of a new vascular sprout (**Figure S1A**). We note that, due to the technical limits, no methods have yet allowed to perform simultaneous assessment of endothelial layer formation, controlled reorganization of a well-defined monolayer boundary, assessment of the cell growth itself at the single-cell level, with sub-cellular resolution of biochemical processes. To address the challenges outlined above, we developed a new platform through the use of optically accessible micro-culture system permitting the following sequential steps: a) generating VEGF gradient in the collagen matrix, b) growth of a fully formed endothelial layer within confined boundaries adjacent to the matrix, c) un-constraining of a part of the monolayer boundary of pre-defined size which enables cells adjacent to this boundary to start invading the adjacent 3D matrix and d) tracking in real time the formation of the resultant sprout, which includes cell migration and the accompanying signaling processes.

To achieve this, we used a polydimethylsiloxane (PDMS) device with two chambers divided by a 50 $\mu$m thin membrane containing 40 $\mu$m wide openings. The device is then bonded onto the glass substratum and was coated with Type 1 rat tail collagen. Details are described in Methods section in the main body of the paper. To generate collagen gel, final concentration of 2.7 mg/ml collagen mixture of DMEM 10x (Sigma, D2429), NaOH 1N, PBS and collagen rat tail 1 (BD Biosciences) was stored in 4 °C fridge for 1 hour, were introduced from upper chamber and incubated at 37 °C for 1 hour in a humid environment for complete polymerization. Surface tension prevents non-polymerized collagen from flooding through openings. Next, endothelial cells of $8 \times 10^6$ cells/ml were induced from lower chamber in confined boundaries and cultured > 15 hours until complete monolayer is formed. The size of the openings was sufficient to contain the cell monolayer and prevent it from invasive spread into the 3D collagen matrix in the absence of the VEGF input. They were also effective in not allowing collagen solution to seep through due to high surface tension. Finally, VEGF was introduced from the upper chamber to create gradient formation for directional motility (**Figure S1B**). This arrangement allowed us to regulate the sprout formation location, treating the membrane with openings as a model of the basement membrane that is partially degraded prior to endothelial sprout formation.

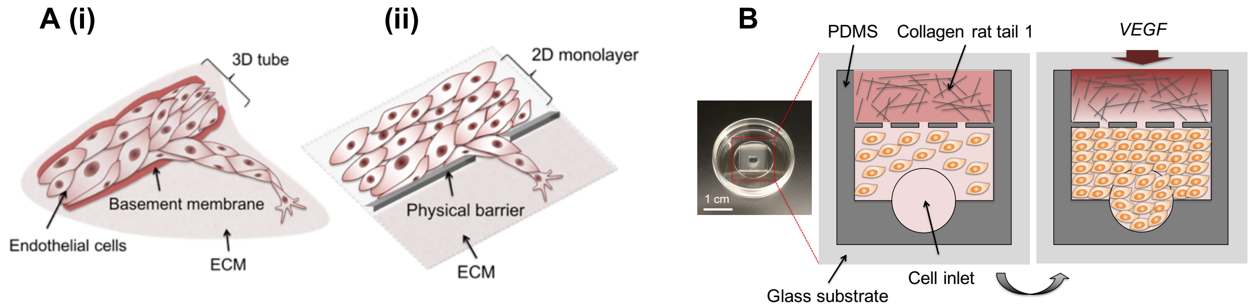


**Figure S1. (A)** Schematics of (i) an *in vivo* angiogenic blood vessels sprouting in the extracellular matrix following a partial local digestion of the basement membrane and (ii) *in vitro* 2D model proposed here with the endothelial cell monolayer mimicking the monolayer of the parental blood vessel, and openings in the barrier mimicking partially digested basement membrane. **(B)** A photograph of the PDMS device on a glass substratum with schematics describing the sequence of experimental procedures: 1) collagen injection and polymerization, followed by cell seeding and monolayer formation in the adjacent chamber (left), and 3) the imposition of the of a VEGF gradient within the collagen matrix promoting directional collective cell invasion and migration into the adjacent matrix-filled chamber. Scale bar: 1 cm.

To analyze the gradient formation inside the matrix, we performed a finite element (COMSOL) simulation with an isotropic, 0 mol/cm$^2$ (bottom) and 1 mol/cm$^2$ (top) as an initial boundary condition with the diffusivity of $0.45 \times 10^{-7}$ cm$^2$/s. The simulation results suggested that a linear gradient inside the matrix was generated from the source to the monolayer boundaries over 24 hours (**Figure S2A**). To confirm the simulation result, we filled the upper chamber with collagen gel (2.7 mg/ml) followed by an addition of FITC tagged dextran (70 kDa) and measured the fluorescence intensity distribution over the length of the collagen layer at 24 hours post dextran application. We also compared the result for diffusion in the chamber without collagen gel. Without collagen matrix, the diffusion profile was noisier and dynamically variant over time, while collagen filled condition produced more consistent spatiotemporal gradient development over time (**Figure S2B**).

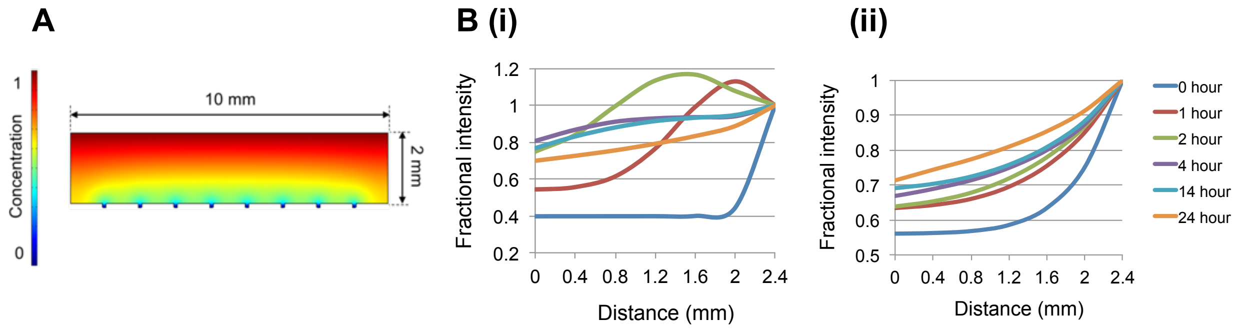


**Figure S2.** Computational and experimental analysis of the imposed VEGF gradient dynamics **(A)** A finite element simulation (using the COMSOL software) of the VEGF gradient dynamics within a model chamber reproducing the experimental geometry of the 3D chamber with the extracellular matrix. Color scale represents relative VEGF concentrations. **(B)** The experimental analysis of the fractional fluorescent intensity of the 70kDa fluorescent dextran was measured across the 3D chamber from the holes to outlets over 24 hours with (i) no collagen and (ii) 2.7 mg/ml collagen conditions.

We observed a substantial difference in the behavior of the cells during cell migration through the narrow openings (holes) to the adjacent chambers that were either not filled or filled with the collagen gel. Whereas the migration that was occurring in 2D on the flat glass led to the collective cell movement, with cells passing through the hole, fanning out, spreading on the 2D surface and forming large lamellipodia (**Figure S3B (i)**), the 3D invasive migration into the collagen filled environment resulted in formation of narrow sprouts with 1-2 leading cells, and much smaller and more elongated cell shapes (**Figure S3B (ii)**). We then explored using this 3D migration model in the collagen gel, whether the cell migration would be accompanied by consistently elevated $Ca^{2+}$ levels. Using MS1 endothelial cells expressing genetically GCaMP5 $Ca^{2+}$ probe, we found that the level of intracellular $Ca^{2+}$ initially increased in multiple cells around the areas adjacent to the opening (**Figure S3C**). This result indicated that the initial reorganization of the layer prior and during initiation of cell invasion can affect groups of cells around the opening, likely due to the joint effects of VEGF input (Noren et al., 2016), mechanical deformation (Hung et al., 2016) and cell-cell communication through gap junctions (Ellison et al., 2016). Again, we found that cells formed well-pronounced sprouts with $Ca^{2+}$ signaling persistently activated at 1-2 'Tip' cells guiding the sprout extension. These results are consistent with our prior results *in vivo*, in zebra fish, suggesting that 'Tip' cells extending into the matrix rich environment have sustained up-regulation of intracellular $Ca^{2+}$ signaling (Noren et al., 2016). These results support the relevance of long term intracellular $Ca^{2+}$ up-regulation for endothelial cells explored in the main body of the paper.

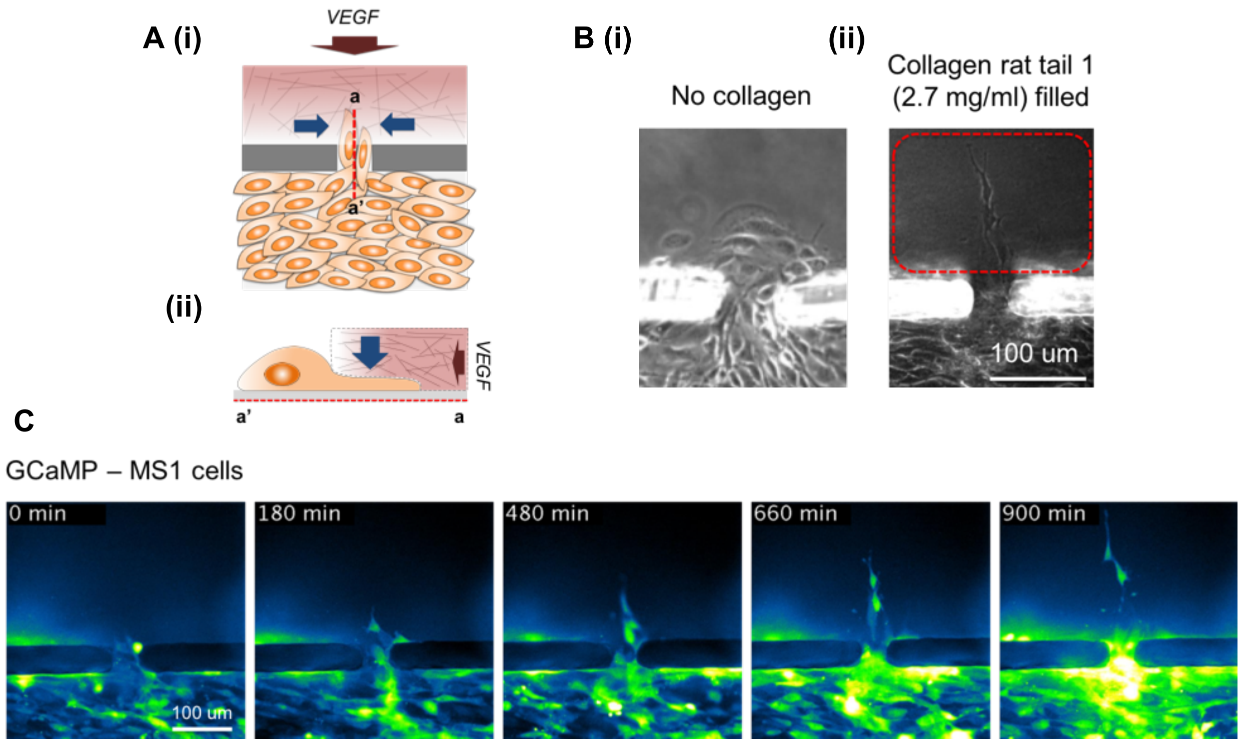


**Figure S3. (A)** Schematics of the cell movement through the holes mimicking areas of degraded basement membrane in response to the VEGF gradient input. The area outside the physical barrier may or may not be filled with collagen matrix (see panel B). A cross section view of a-a' in A (i) is shown in A (ii). **(B)** Representative phase contrast images of MS1 cells invading through a small hole (width of 40 μm) with (i) no collagen and (ii) collagen gel in the adjacent chamber filled condition. Red dotted box indicates the region of collagen matrix. Scale bar: 100 μm. **(C)** Representative time-lapse fluorescent images of GCaMP5 expressing MS1 cells invading collagen matrix. Scale bar: 100 μm.

### References

Ellison, D., Mugler, A., Brennan, M.D., Lee, S.H., Huebner, R.J., Shamir, E.R., Woo, L.A., Kim, J., Amar, P., Nemenman, I.*, et al.* (2016). Cell-cell communication enhances the capacity of cell ensembles to sense shallow gradients during morphogenesis. Proc Natl Acad Sci U S A *113*, E679-688.

Hung, W.C., Yang, J.R., Yankaskas, C.L., Wong, B.S., Wu, P.H., Pardo-Pastor, C., Serra, S.A., Chiang, M.J., Gu, Z.Z., Wirtz, D.*, et al.* (2016). Confinement Sensing and Signal Optimization via Piezo1/PKA and Myosin II Pathways. Cell Rep *15*, 1430-1441.

Noren, D.P., Chou, W.H., Lee, S.H., Qutub, A.A., Warmflash, A., Wagner, D.S., Popel, A.S., and Levchenko, A. (2016). Endothelial cells decode VEGF-mediated $Ca^{2+}$ signaling patterns to produce distinct functional responses. Science Signaling *9*, ra20.
