## Supplementary Methods S2 for "A molecular clock controls periodically driven cell migration in confined spaces"

### Methods S2. Peak detection analysis

We conducted the peak detection analysis and put the results for the experiments below. Details of the analysis are described in Methods (Image analysis section) in the main body of the paper. The numbers refer to the peaks detected by the algorithm. The figures labelled to refer to the data in the main and supplementary data.

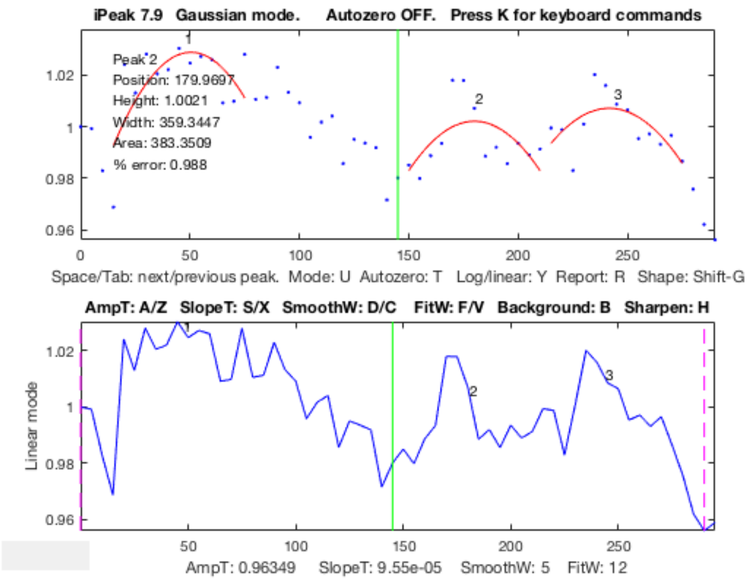


**(A)** Supplementary Figure 1C

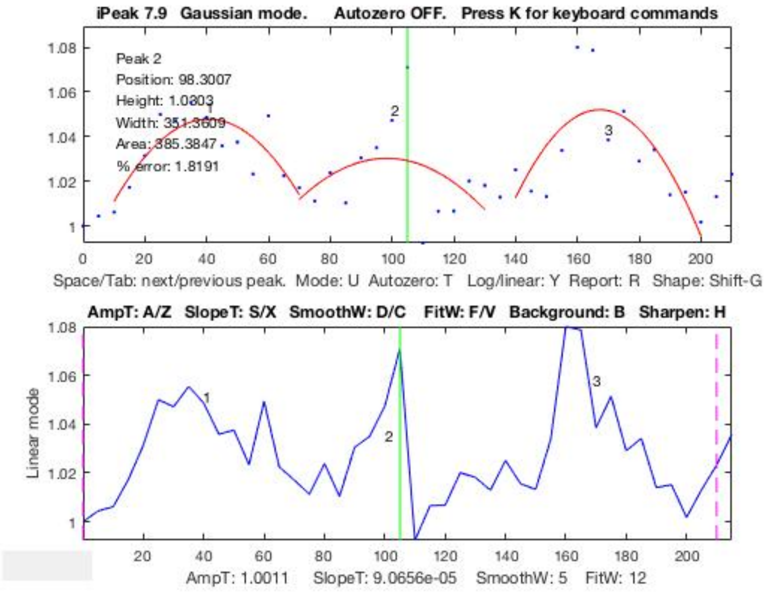


**(B)** Figure 1B (ii) Control conditions

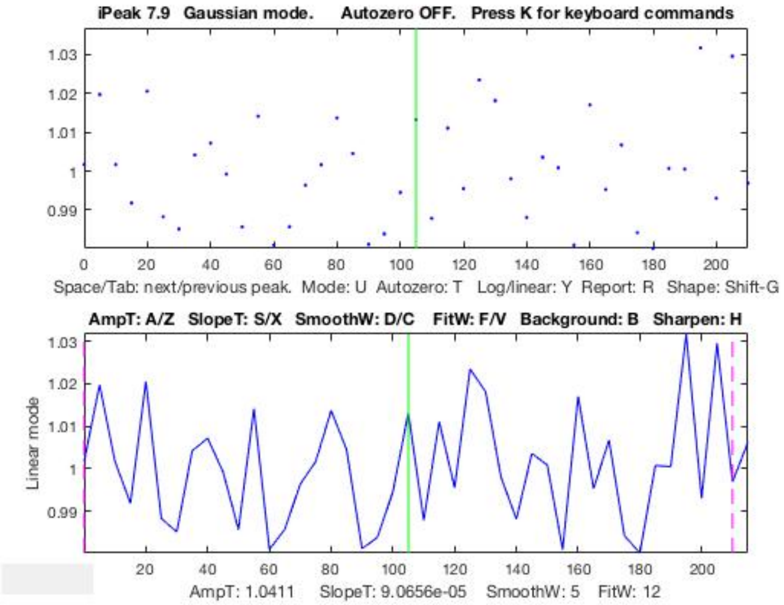


**(C)** Figure 1B (ii) BAPTA treatment

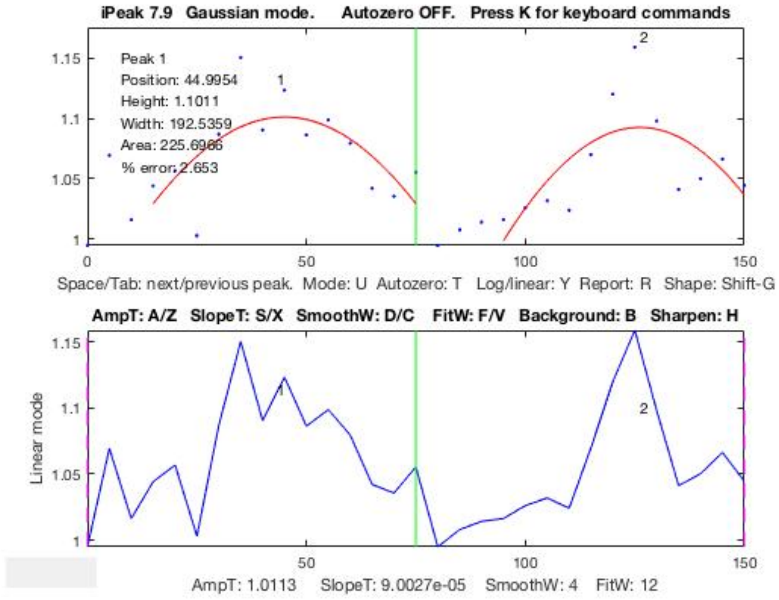


**(D)** Figure 1C (ii) RhoA activity analysis

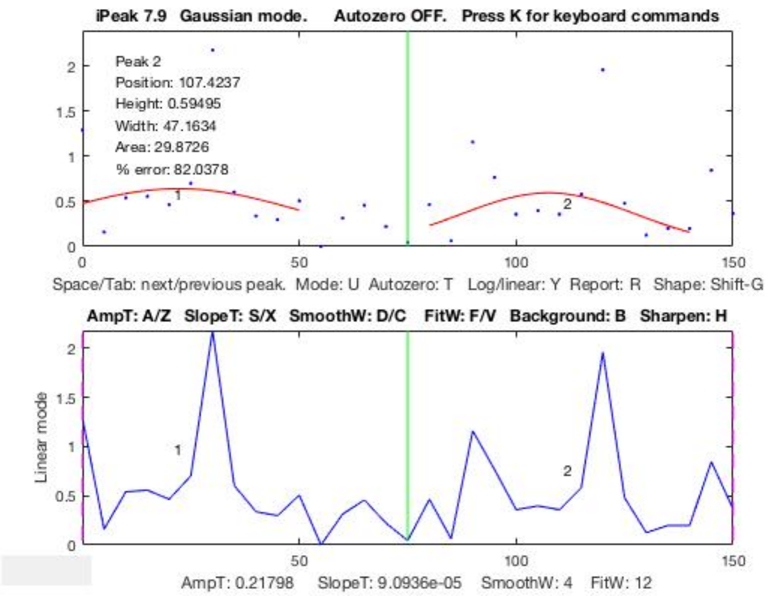


**(E)** Figure 1C (ii) Migration speed

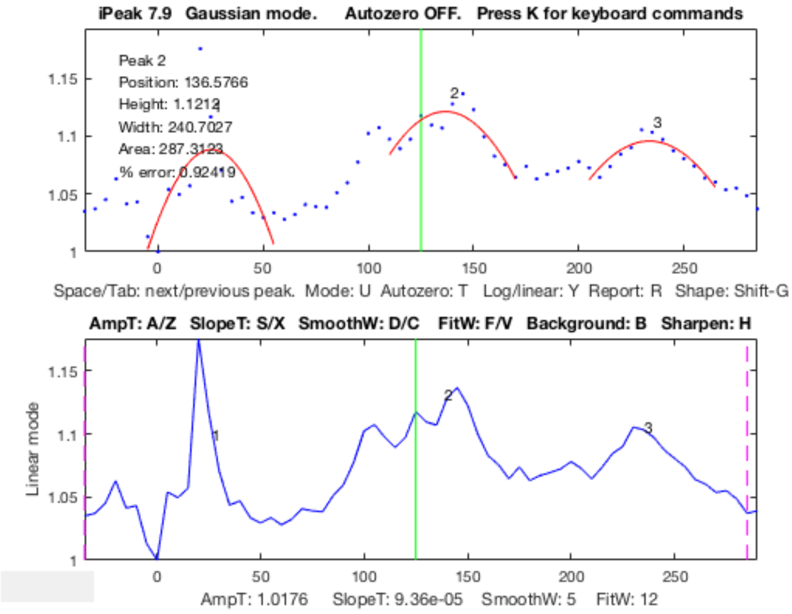


**(F)** Figure 2A (iii)

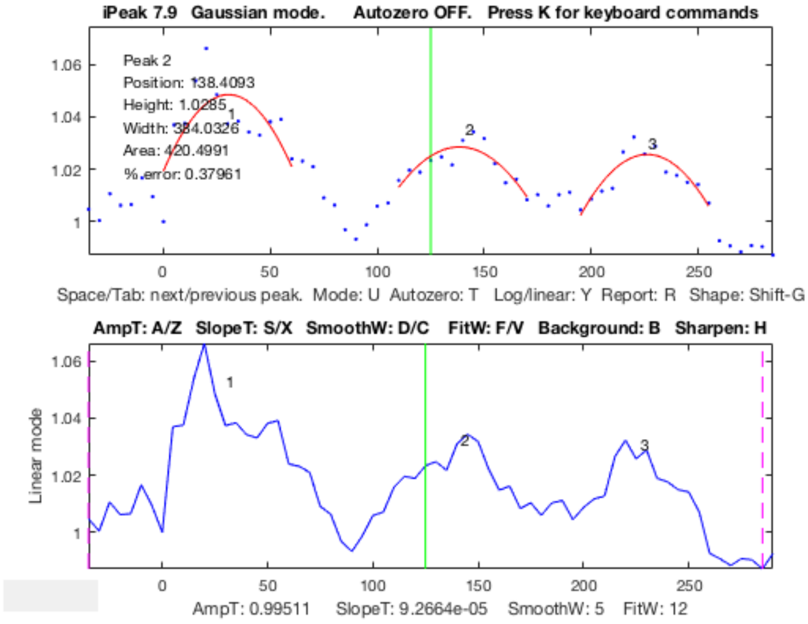


**(G)** Supplementary Figure 2A

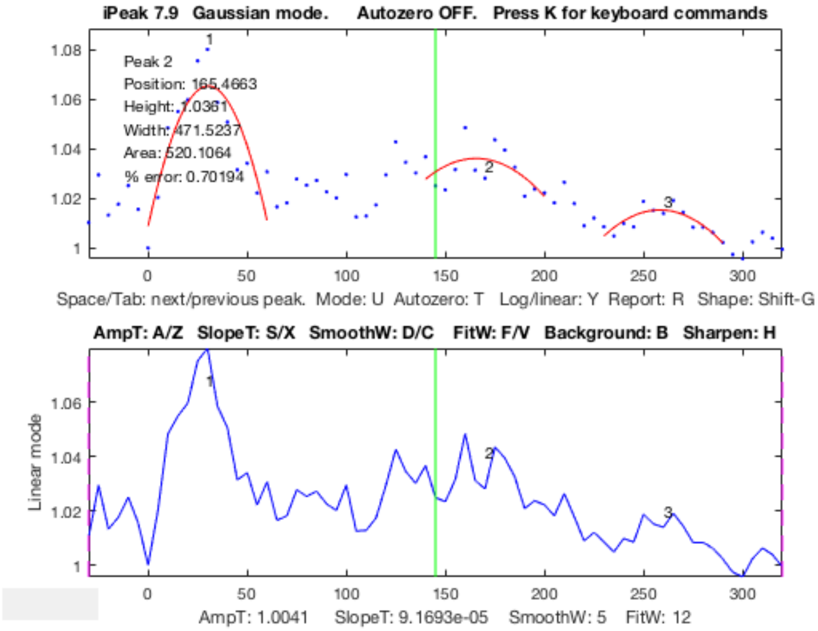


**(H)** Supplementary Figure 2B (i)

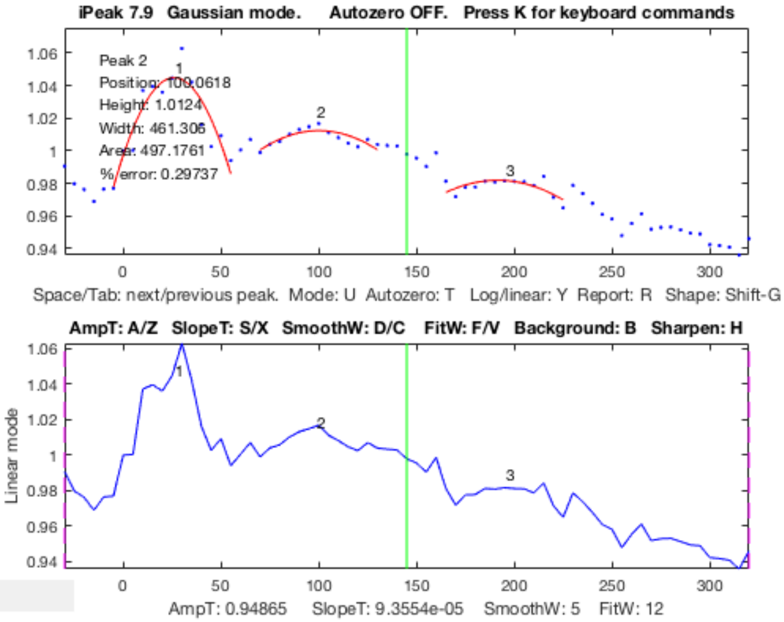


**(I)** Supplementary Figure 2B (ii)

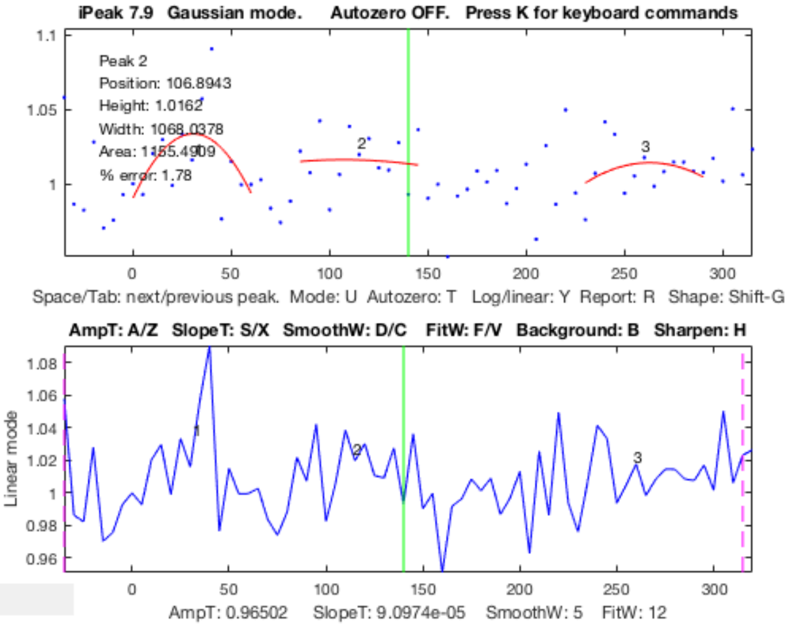


**(J)** Supplementary Figure 2C (i)

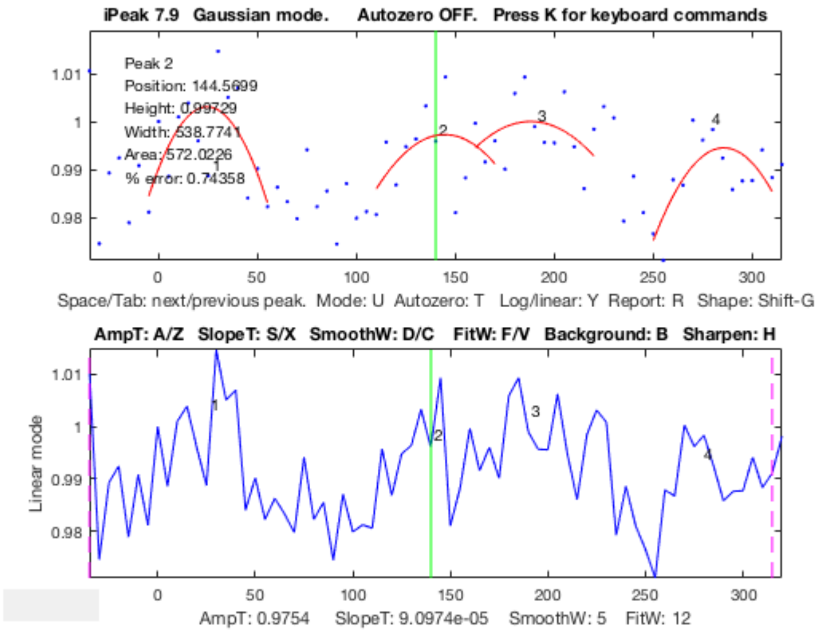


**(K)** Supplementary Figure 2C (ii)
