## Supplementary Methods S3 for "A molecular clock controls periodically driven cell migration in confined spaces"

### Methods S3.
### Mathematical model of $Ca^{2+}$-induced oscillatory RhoA dynamics

#### 1. Model Overview

We modeled the biochemical circuit putatively underlying the molecular clock described in the main text as a set of ordinary differential equations (ODEs), describing the dynamics of concentrations of various biomolecular species. Briefly, in accordance with the results described in the manuscript, we assumed that the transition from MT polymerization to MT depolymerization promotes GEF-H1 activation and has a negative effect on GEF-H1 abundance, which triggers a negative feedback resulting in an increase of abundance of mDia1, and elevated actin and MT polymerization.  Furthermore, active GEF-H1 can trigger mDia1 activation, due its function as a RhoA-GEF, triggering transitions between RhoA-GTP and RhoA-GDP complexes. Moreover, downregulation of the GEF-H1 abundance in turn upregulates mDia1 concentration, leading to the second GEF-H1-mediated negative feedback loop (**Figure M1**).

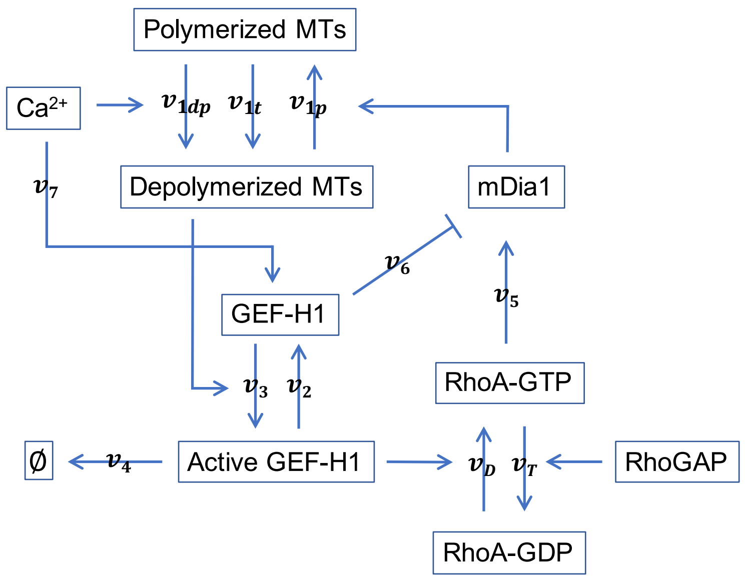


**Figure M1. Schematic of the circuit driving the molecular clock leading to oscillation in RhoA activity.** The arrows indicate biochemical activation (pointed) or inhibition (blunted), with the letters referring to the biochemical reaction rates, as described below.

The ODEs were based on saturable linear kinetics, recognizing that this may over-simplify some of the non-linear kinetic mechanisms. This assumption allowed us to minimize the number of the parameters required to describe the system kinetics. A posteriori, linear saturable kinetics was sufficient to describe the dynamics of the biochemical processes assayed in the manuscript, justifying the relatively simple setup (see the discussion below). In the schematic above and equations below, [*MT pol*] refers to the concentration of tubulin in polymerized microtubules, whereas [*MT depol*] is the concentration of depolymerized tubulin subunits. [*GEFH1*] is the total concentration of GEF-H1, whereas [*Active GEFH1*] is the concentration of GEF-H1 unbound with microtubules. [*RhoA GDP*] and [*RhoA GTP*] are the concentrations of RhoA GDP and RhoA GTP respectively. [*RhoGAP*] is the concentration of RhoGAP and [*mDia1*] is the concentration of mDia1. [$Ca^{2+}$] is the concentration of intracellular $Ca^{2+}$.

$$v_{1t} = \frac{k_{1t} \cdot [MT\ pol]}{K_{1t} + [MT\ pol]} \quad \text{(R1)}$$

is the rate of microtubule depolymerization under the basal conditions.

$$v_{1dp} = \frac{k_{1dp} \cdot [Ca^{2+}] \cdot [MT\ pol]}{K_{1dp} + [MT\ pol]} \quad \text{(R2)}$$

is the rate of microtubule depolymerization under the elevated $Ca^{2+}$ conditions.

$$v_{1p} = \frac{k_{1p} \cdot [mDia1] \cdot [MT\ depol]}{K_{1p} + [MT\ depol]} \quad \text{(R3)}$$

is the rate of mDia1-dependent microtubule polymerization.

$$v_2 = \frac{k_2 \cdot [Active\ GEFH1]}{K_2 + [Active\ GEFH1]} \quad \text{(R4)}$$

is the rate of constitutive GEF-H1 inactivation.

$$v_3 = \frac{k_3 \cdot [MT\ depol] \cdot [GEFH1]}{K_3 + [GEFH1]} \quad \text{(R5)}$$

is the rate of GEF-H1 activation mediated by unbinding from dissociating MTs.

$$v_4 = k_4 \cdot [Active\ GEFH1] \quad \text{(R6)}$$

is the rate of constitutive degradation of free GEF-H1.

$$v_D = \frac{k_D \cdot [Active\ GEFH1] \cdot [RhoA\ GDP]}{K_D + [RhoA\ GDP]} \quad \text{(R7)}$$

is the rate of RhoA activation mediated by active GEF-H1.

$$v_T = \frac{k_T \cdot [RhoGAP] \cdot [RhoA\ GTP]}{K_T + [RhoA\ GTP]} \quad \text{(R8)}$$

is the rate of RhoA inactivation mediated by a RhoA GAP protein.

$$v_5 = \frac{k_5 \cdot [RhoA\ GTP]}{K_5 + [RhoA\ GTP]} \quad \text{(R9)}$$

is the rate of mDia1 activation downstream of active RhoA.

$$v_6 = \frac{k_6 \cdot [GEFH1]}{K_6 + [GEFH1]} \quad \text{(R10)}$$

is the rate of GEF-H1 dependent decrease in active mDia1 (due to decreasing mDia1 abundance).

$$v_7 = \frac{k_7 \cdot [Ca^{2+}]}{K_7 + [Ca^{2+}]} \quad \text{(R11)}$$

is the rate of $Ca^{2+}$-dependent GEF-H1 synthesis.

The model was implemented with the following ODE equations based on the reaction rates (R1-R11) described above. Parameters used in the equations are listed in **Table M1** below. We assumed that the rate of intracellular $Ca^{2+}$ increase (the derivative rather the concentration itself) exponentially decays in the system after initial treatment. The ODEs were simulated using custom written code in MATLAB 2018:

$$\frac{d[Ca^{2+}]}{dt} = a \cdot e^{-\gamma t} \quad \text{(E1)}$$

$$\frac{d[MT\ depol]}{dt} = v_{1dp} + v_{1t} - v_{1p} \quad \text{(E2)}$$

$$\frac{d[MT\ pol]}{dt} = v_{1p} - v_{1dp} - v_{1t} \quad \text{(E3)}$$

$$\frac{d[GEFH1]}{dt} = v_2 - v_3 + v_7 - v_4 \quad \text{(E4)}$$

$$\frac{d[Active\ GEFH1]}{dt} = v_3 - v_2 \quad \text{(E5)}$$

$$\frac{d[RhoA\ GDP]}{dt} = v_T - v_D \quad \text{(E6)}$$

$$\frac{d[RhoA\ GTP]}{dt} = v_D - v_T \quad \text{(E7)}$$

$$\frac{d[mDia1]}{dt} = v_5 - v_6 \quad \text{(E8)}$$

Model parameters, given in the Table below were either taken from the indicated reference or estimated through data fitting as explained below.

**Table M1. Parameters used in the equations (R1-R11, E1-E8) (units in parentheses)**

| Parameter | Value ($k$'s: min$^{-1}$, $K$'s: nM) | Reference |
|---|---|---|
| $\alpha$ | 0 (basal), 10000 ($Ca^{2+}$) | Assumed |
| $\gamma$ | 10 | Assumed |
| $k_{1dp}$ | 0.000475 | Estimated |
| $K_{1dp}$ | 690 | Estimated |
| $k_{1t}$ | 5 (basal), 0 ($Ca^{2+}$) | Estimated |
| $K_{1t}$ | 1200 | Estimated |
| $k_{1p}$ | 2.4 | Estimated |
| $K_{1p}$ | 5.6 | Estimated |
| $k_2$ | 25.25 | Estimated |
| $K_2$ | 20 | (Byrne et al., 2016) |
| $k_3$ | 2.525 | Estimated |
| $K_3$ | 30 | (Byrne et al., 2016) |
| $k_4$ | 0.00001 | Estimated |
| $k_5$ | 0.5 | Estimated |
| $K_5$ | 40 | Estimated |
| $k_6$ | 0.18 | Estimated |
| $K_6$ | 105 | Estimated |
| $k_7$ | 0.0000135 | Estimated |
| $K_7$ | 26 | Estimated |
| $k_D$ | 4.5 | Estimated |
| $K_D$ | 170 | (Byrne et al., 2016) |
| $k_T$ | 0.3 | Estimated |
| $K_T$ | 10 | (Byrne et al., 2016) |
| *[RhoGAP] (nM)* | 40 | Estimated |

**Basal Condition vs. High $Ca^{2+}$ Conditions**
The key trigger of the molecular clock is the persistent increase in the intracellular $Ca^{2+}$ levels. Prior to this input, the system is in a basal steady state. We therefore, pre-equilibrated the model prior to the input addition, using the following initial conditions and ensuring that the model reached the steady state (all units are nM). (**Table M2, Figure M2A**):

**Table M2. Parameters used before training the model.**

| Parameter | Value ($k_d$: min$^{-1}$, $K_d$: nM) |
|---|---|
| $k_{1dp}$ | 0.0005 |

| | |
|---|---|
| $K_{1dp}$ | 600 |
| $k_{1p}$ | 3 |
| $K_{1p}$ | 10 |
| $k_6$ | 0.2 |
| $K_6$ | 100 |
| $k_7$ | 0.000015 |
| $K_7$ | 20 |

We found that the system reached the steady state at approximately 10 hours, reading the following values (all units are nM):

[$Ca^{2+}$] = 0, [*MT pol*] = 46.2, [*MT depol*] =3.8, [*GEFH1*] = 8.21, [*Active GEFH1*] = 1.78, [*RhoA GDP*] = 28.93, [*RhoA GTP*] = 1.07 and [*mDia1*] = 0.19

As noted above, we assumed that intracellular $Ca^{2+}$ increases exponentially and maintains at concentration level (1 $\mu$M) according to the equation E1 (**Figure M2C**). We found through simulations that model can robustly generate oscillatory responses within specific parameter range, with the equations we have postulated, with one exception. The initial concentration of mDia1 at the time of $Ca^{2+}$ input needed to be reset to low level (assumed to be zero) to allow for the oscillatory response, suggesting that the sensitivity of MT polymerization to mDia1 levels was not linear. With these assumptions, the equilibrated initial concentrations and parameters described above, the model displayed oscillations with 3 peaks over 5-hour period of time, matching our experimental results (**Figure M2B and Figure 2B**).

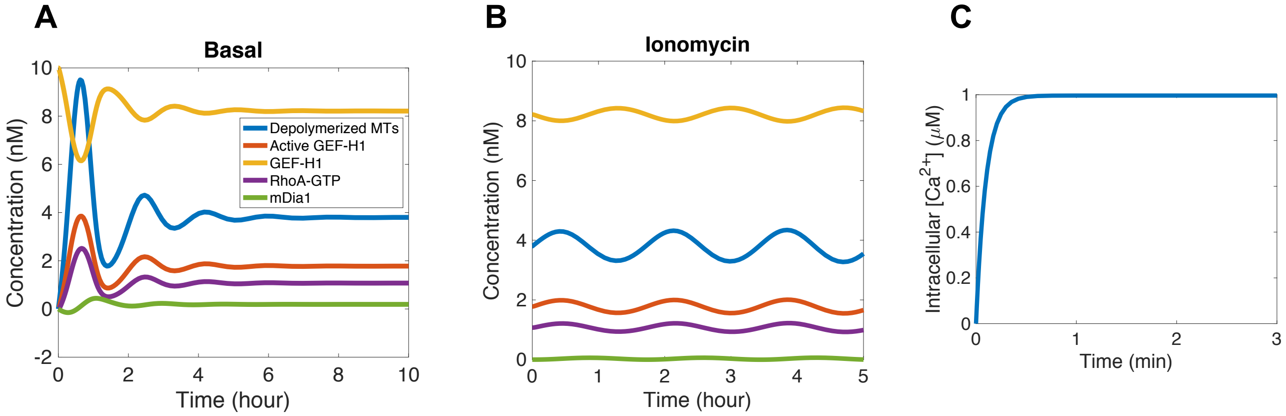


**Figure M2. Modeling results** of the dynamics of depolymerized MTs (blue), active GEF-H1 (red), GEF-H1 (yellow), RhoA-GTP (purple) and mDia1 (green) in (A) basal state equilibrations and (B) ionomycin treatment cases. (C) Model assumption regarding intracellular [$Ca^{2+}$] level increase.

#### 2. Model Training

The model was parameterized by comparing the simulation output to a subset of experimental data. In particular, the model results of the active GEF-H1 dynamics from **Figure M2B** were compared with the mean value of active GEF-H1 in **Figure 2B** acquired from multiple sets of experiments. We chose results of active GEF-H1 because it shows oscillations ubiquitously across different types of cells including HUVECs, MS1 and A375 cells. In the training, we minimized the following metric:

$$\sum_{Time} \left(Active\ GEFH1\ (model) - Folds\ of\ active\ GEFH1\ over\ total\ GEFH1\ (experiment)\right)^2$$

Based on the Direct algorithm method used from previous studies (Jones et al., 1993; Vanaja et al., 2018), we locally varied parameters +/- 30 % with 5 % increments and calculated the metric above to find the parameter generating the minimized output. The fit was particularly sensitive to the following rates: MT polymerization ($v_{1p}$) and depolymerization ($v_{1dp}$), the $Ca^{2+}$ dependent GEF-H1 synthesis ($v_7$) and the GEF-H1-dependent of inhibition of mDia1 abundance ($v_6$), whereas variation of other parameters did not result in a substantial change in the metric. The initial values of the parameters were taken whenever possible from (Cytrynbaum et al., 2004; Byrne et al., 2016; Sha et al., 2018; Floyd et al., 2017; Jilkine et al., 2007). With the adjusted parameters (**Table M1**), we were able to generate oscillated dynamics of the system reflecting more closely to the experimental results of active GEF-H1 (**Figure M2**). We further explored the parameter sensitivity of the model, as described in the section below.

##### 3. Sensitivity Analysis

To understand what parameters are more critical to determine the frequency and decreasing/increasing amplitude of the oscillations, we conducted the parameter sensitivity analysis. We varied each parameter +/- 60 % from its nominal value and measured the relative changes in the RhoA oscillation frequency and amplitude over a simulated time of 50 hours. The frequency was measured using the automated peak detection method (Methods S2 and the Methods section of the main body of the paper) with the following threshold values: amplitude threshold 0.01, slope threshold 0.0001, smooth width 4 and fit width 10. The change in the amplitude was evaluated as the ratio of the amplitudes of the first and the last peaks detected in a range of 50 hours (**Figure M3**).

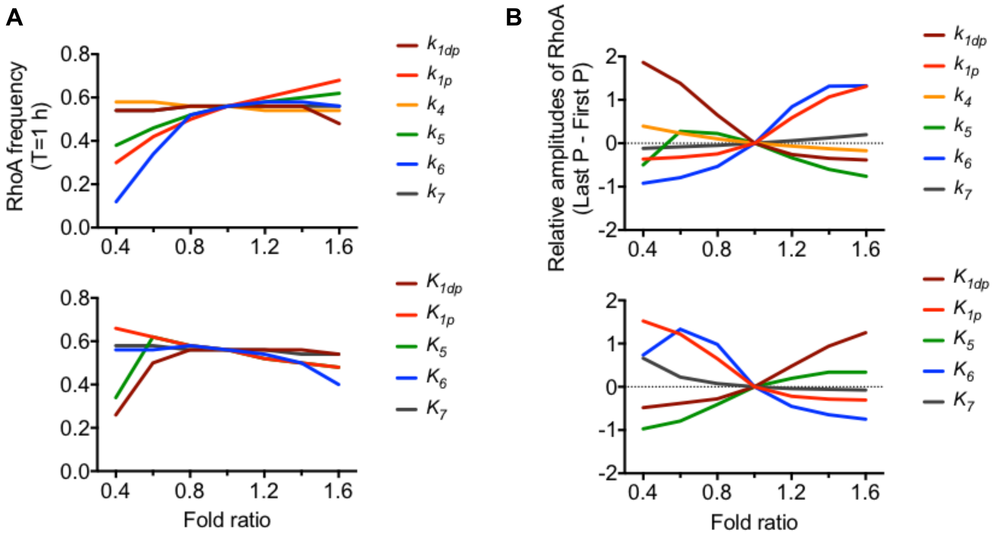


**Figure M3. Sensitivity analysis of the model** (A) RhoA frequency (number of pulses per 1 hour) and (B) Relative amplitudes of RhoA at the last vs. first time point. See text for further details.

Based on the analysis results, MT polymerization parameters ($k_{1p}$, $K_{1p}$) approximately linearly influenced the RhoA oscillation frequency. In particular, RhoA oscillated with higher frequencies when MTs polymerization occurred faster with lower $K_d$. MT depolymerization rate decreased the RhoA frequency, especially when $K_d$ constant was lower than the nominal value. The frequency was also very sensitive to the variation of the values of the parameters determining the rate of GEF-H1-dependent mDia1 inactivation ($k_6$, $K_6$). The frequency was reduced as $k_6$ was reduced and $K_6$ was increased, indicating that RhoA oscillations slow down either when the rate of mDia inhibition is reduced (low $k_6$) or when the inhibition reaches the maximum rate slower than normal condition (high $K_6$). The parameters controlling both MT depolymerization and polymerization rates also determined whether the amplitude of the oscillations gradually increased or decreased over the course of the simulations. For example, the degree of the amplitude decay was enhanced when MTs depolymerized faster ($k_{1dp}$) with low $K_{1dp}$, whereas the degree of the amplitudes enhancement was increased for faster rates ($k_{1p}$) of MT polymerization with low $K_{1p}$. The values of the parameters determining the rate of GEF-H1-dependent mDia1 inactivation were also found to control the amplitude change, particularly with the increasing $k_6$ and decreasing $K_6$ leading to a strong amplitude amplification.

#### 4. Model Perturbations to test the proposed mechanisms underlying the molecular clock

As described in the main text, the model was used in testing of the proposed mechanisms of the molecular clock driving cell migration in physically confining spaces. The ultimate test was the analysis of whether the frequency and amplitude of the RhoA oscillation and cell migration could be controlled in the predictive fashion by perturbations of the clock components. To model various pharmacological perturbations, as described below, we varied rates of the indicated reactions. Since we did not know the exact effect that a drug may have on the reaction parameters, we tested two values for the reactions rates, while also varying the corresponding saturation constants (K's) by 20 %, 0 %, +20 % of their nominal values. We thus obtained 6 predicted dynamic behaviors, as shown in Fig. 6 of the main text (highlighting the one most closely agreeing with the corresponding experimental observation). We observed that the level of MT acetylation was enhanced following the treatment of C646, indicative of an increase of MT polymerization (**Supplementary Figures 7E**). Also, GEF-H1 synthesis was inhibited following the cell treatment with C646 under high intracellular $Ca^{2+}$ condition, as this drug also targets the binding of CREB binding protein to p300 necessary for transcriptional activation. Thus, we varied the parameters regulating the MT polymerization ($k_{1p}$, $K_{1p}$) and GEF-H1 synthesis rate ($k_7$, $K_7$), under the assumption that the C646 treatment results in an increased MT polymerization rate and a reduced GEF-H1 synthesis rate vs ionomycin treatment alone. We thus increased the value of $k_{1p}$ by 10 % and 200 % and decreased the value of $k_7$ by 40% and 90% (in this case $K_7$ was varied by 30% rather the usual 20%). To account for mDia1 perturbation by SMIFH2, we assumed that the effect will be to inhibit MT polymerization (Bartolini et al., 2016; Palazzo et al., 2001). Therefore, we varied $k_{1p}$ by 3% and 30%. In the case of MT targeting agents, we assumed that nocodazole treatment leads to increased MT depolymerization rate. Therefore, we varied $k_{1t}$ by 10 % and 200 %. By contrast, we hypothesized that paclitaxel treatment induces more rapid MT polymerization rate. Thus, we increased the MT polymerization rate, $k_{1p}$, by 10 % and 200 %. (**Table M3**) shows the parameter values producing the best agreement with the experimental data.

**Table M3. Parameters in perturbed conditions**

| Condition | Parameter | Value (*k's*: min$^{-1}$, *K's*: nM) |
|---|---|---|
| C646 + Ionomycin | $k_{1p}$ | 4.8 |
| | $K_{1p}$ | 4.48 |
| | $k_7$ | 0.0000125 |
| | $K_7$ | 26 |
| SMIFH2 + Ionomycin | $k_{1p}$ | 0.072 |
| | $K_{1p}$ | 6.72 |
| Nocodazole | $k_{1t}$ | 5.5 |
| | $K_{1t}$ | 1440 |
| Paclitaxel | $k_{1p}$ | 2.64 |
| | $K_{1p}$ | 6.72 |

#### References

Bartolini, F., Andres-Delgado, L., Qu, X.Y., Nik, S., Ramalingam, N., Kremer, L., Alonso, M.A., and Gundersen, G.G. (2016). An mDia1-INF2 formin activation cascade facilitated by IQGAP1 regulates stable microtubules in migrating cells. Mol Biol Cell *27*, 1797-1808.
Byrne, K.M., Monsefi, N., Dawson, J.C., Degasperi, A., Bukowski-Wills, J.C., Volinsky, N., Dobrzynski, M., Birtwistle, M.R., Tsyganov, M.A., Kiyatkin, A.*, et al.* (2016). Bistability in the Rac1, PAK, and RhoA Signaling Network Drives Actin Cytoskeleton Dynamics and Cell Motility Switches. Cell Syst *2*, 38-48.
Cytrynbaum, E.N., Rodionov, V., and Mogilner, A. (2004). Computational model of dynein-dependent self-organization of microtubule asters. J Cell Sci *117*, 1381-1397.
Floyd, C., Jarzynski, C., and Papoian, G. (2017). Low-dimensional maniford of actin polymerization dynamics. New J Phys *19*, 125012.
Jilkine, A., Maree, A.F., and Edelstein-Keshet, L. (2007). Mathematical model for spatial segregation of the Rho-family GTPases based on inhibitory crosstalk. Bull Math Biol *69*, 1943-1978.
Jones, D.R., Perttunen, C.D., and Stuckman, B.E. (1993). Lipschitzian Optimization without the Lipschitz Constant. J Optimiz Theory App *79*, 157-181.
Palazzo, A.F., Cook, T.A., Alberts, A.S., and Gundersen, G.G. (2001). mDia mediates Rho-regulated formation and orientation of stable microtubules. Nat Cell Biol *3*, 723-729.
Sha, Z., Zhao, J., and Goldberg, A.L. (2018). Measuring the Overall Rate of Protein Breakdown in Cells and the Contributions of the Ubiquitin-Proteasome and Autophagy-Lysosomal Pathways. Methods Mol Biol *1844*, 261-276.
Vanaja, K.G., Timp, W., Feinberg, A.P., and Levchenko, A. (2018). A Loss of Epigenetic Control Can Promote Cell Death through Reversing the Balance of Pathways in a Signaling Network. Mol Cell *72*, 60-70.
